# Integrating proteomic data with metabolic modelling provides insight into key pathways of *Bordetella pertussis* biofilms

**DOI:** 10.1101/2022.05.27.493021

**Authors:** Hiroki Suyama, Laurence Don Wai Luu, Ling Zhong, Mark J. Raftery, Ruiting Lan

## Abstract

Pertussis, commonly known as whooping cough is a severe respiratory disease caused by the bacterium, *Bordetella pertussis*. Despite widespread vaccination, pertussis resurgence has been observed globally. The development of the current acellular vaccine (ACV) has been based on planktonic studies. However, recent studies have shown that *B. pertussis* readily forms biofilms. A better understanding of *B. pertussis* biofilms is important for developing novel vaccines that can target all aspects of *B. pertussis* infection. This study compared the proteomic expression of biofilm and planktonic *B. pertussis* cells to identify key changes between the conditions. Major differences were identified in virulence factors including an upregulation of toxins (adenylate cyclase toxin and dermonecrotic toxin) and downregulation of pertactin and type III secretion system proteins in biofilm cells. To further dissect metabolic pathways that are altered during the biofilm lifestyle, the proteomic data was then incorporated into a genome scale metabolic model using the integrative metabolic analysis tool (iMAT). The analysis revealed that planktonic cells utilised the glyoxylate shunt while biofilm cells completed the full tricarboxylic acid cycle. Differences in processing aspartate, arginine and alanine were identified as well as unique export of valine out of biofilm cells which may have a role in inter-bacterial communication and regulation. Finally, increased polyhydroxybutyrate accumulation and superoxide dismutase activity in biofilm cells may contribute to increased persistence during infection. Taken together, this study modelled major proteomic and metabolic changes that occur in biofilm cells which helps lay the groundwork for further understanding *B. pertussis* pathogenesis.

## Introduction

Whooping cough is a re-emerging severe respiratory disease caused by *Bordetella pertussis*. Following the change from the whole cell vaccine (WCV) to the acellular vaccine (ACV) in many developed countries, there has been an increase in the incidence of whooping cough (1–4). Although most likely multifaceted, waning immunity of the ACV and vaccine driven selection of non-ACV genotypes or strains not expressing one of the ACV antigens have been previously reported as major factors contributing to the re-emergence of pertussis (5, 6). It is evident that an improved ACV is needed to control the infections.

Recent studies have shown that *B. pertussis* readily forms biofilms *in vivo* (7–13). Development of the vaccine has been based on planktonic studies and may not be entirely representative of the infection cycle. Although proteomic comparisons have been performed between biofilm and planktonic *B. pertussis* cells (14–18), little is known about the metabolic reactions that are altered while in the biofilm state. In response to changes in environment, the most widely studied regulator of gene expression in *B. pertussis* is the *Bordetella* virulence gene (Bvg) system (19). The Bvg system controls the expression of most of the virulence factors in *B. pertussis* but has also been implicated in the regulation of metabolism (20). The Bvg system exists in 3 states, Bvg^+^, Bvg^-^ and Bvg^i^ where the expression of virulence genes is active, inactive or intermediately expressed, respectively. There are conflicting studies surrounding the role of the Bvg system in the biofilm process, but studies have linked *Bordetella* biofilm with the Bvg^i^ phase (14, 17, 18, 21–23). Further studies identifying key changes in protein expression and metabolism in biofilm cells can provide an insight into the capabilities of the pathogen and help with understanding the role of biofilms in *B. pertussis* pathogenesis.

Genome scale metabolic models (GSMM) have emerged as a powerful tool in understanding the metabolic capabilities of an organism. The creation of a GSMM begins as a draft network based on annotated enzyme data and the genome. This yields a model with a network of reactions and metabolites within a mathematical matrix. The movement of metabolites through the network is defined as flux i.e., the rate at which the metabolites are consumed or produced. Based on stoichiometric and thermodynamic constraints, permissible minimum and maximum flux values for each reaction can be calculated (24).

GSMMs have been successfully utilised in several pathogens such as *Salmonella enterica* serovar Typhimurium (25)*, Listeria monocytogenes* (26, 27), *Staphylococcus aureus* (28) and *Mycobacterium tuberculosis* (29) to predict important pathways for virulence and growth. Currently, there have been over 6,000 organisms that have been metabolically reconstructed either manually or automatically (30). There have been two GSMMs extensively curated for *B. pertussis* (31, 32). These models showed the metabolic versatility of the organism by identifying minimal nutrient requirements. Additionally, both models were utilised to validate key pathways essential for infection (33). The major drawback of these models is the assumption that all protein products are simultaneously expressed. Many physical and chemical constraints make this assumption unlikely to be true. Additionally, the different conditions in which the organism is grown strongly affects the metabolic processes (34). To predict metabolic reactions reflecting a specific phenotype, such as biofilms, a context specific model should be created. This can be done through the integration of ‘omics’ expression data into the GSMM to enrich pathways reflective of the context (i.e., biofilm). The Integrative Metabolic Analysis Tool (iMAT) (35) is a method that has been developed to incorporate expression data within a metabolic model. The iMAT algorithm uses a mixed integer linear programming (MILP) problem to enrich pathways based on the expression data while maintaining a steady flux distribution and the stoichiometric and thermodynamic constraints. The iMAT model not only creates a phenotype specific model but post-transcriptional regulation or inaccuracies in expression due to experimental limitations can be predicted through this model as the surrounding fluxes would indicate the relative activity of an enzyme in a pathway (35). By comparing the flux distributions of two context specific models, changes in metabolic reactions between phenotypes can be predicted (36).

In this study, proteomic expression data was used to compare biofilm and planktonic *B. pertussis* cells from a representative current circulating strain. Furthermore, the protein expression data was incorporated into a GSMM to create context specific iMAT models to elucidate key metabolic changes that may allow biofilm cells to persist in the host.

## Methods

### Bacterial strains and biofilm growth

A clinical *Bordetella pertussis* strain, L1423 isolated from the 2008-2012 Australian epidemic with genotype *ptxP3/ptxA1/fim3A/prn2* and expressing pertactin, was used as a representative of the predominant cluster I strains (37). The genome has been previously sequenced, and the strain has been used in two separate infection studies in mice (38, 39). *B. pertussis* cells were grown using a previously established method and proteins were extracted (40). Briefly, the *B. pertussis* strain was grown on Bordet-Gengou agar (BG, BD Scientific) for 3-5 days at 37°C. A loopful of pure Bvg^+^ colonies were suspended into Thalen-Ijssel (THIJS) media (41) supplemented with 1% heptakis ((2,6□O□dimethyl) β □ cyclodextrin) and 1% THIJS supplement and grown for 24 h shaking at 180 rpm at 37°C. For planktonic growth, the OD_600_ of the starter culture was adjusted to 0.05/mL in 20 mL THIJS and incubated for 12 h, shaking at 180 rpm (reaching log phase for L1423 (42)). For biofilms, the OD_600_ was adjusted to 0.1/mL and 1 mL of this adjusted culture was seeded into each well of a 24 well plate (15, 43). The 24 well plate was incubated statically for 5 h at 37°C for attachment of cells before the media was refreshed to remove non-adherent cells (14, 18). After 96 h of incubation under agitation (60 rpm), the wells were washed with PBS and then the plate was water bath sonicated at 37 kHz for 2 min to detach cells (44, 45). The planktonic and biofilm cells were then probe sonicated and proteins extracted as described by Luu *et al*. (40). Six biological replicates per condition were performed.

### Confocal laser scanning microscopy analysis

To confirm biofilm maturity, confocal laser scanning microscopy (CLSM) was used. This method was adapted from Cattelan *et al*. (46). The *B. pertussis* cells were grown on a glass coverslip angled at 45° in the same method as described above. The biofilm was imaged at 24, 48, 72 and 96 h. The biofilm was fixed with 4% paraformaldehyde and stained with SYTO 9 (Thermo Fisher Scientific) fluorescent dye. The coverslips were imaged on the FluoView FV1200 inverted confocal microscope (Olympus Life Sciences) at the UNSW Katharina Gaus Light Microscopy Facility (KG-LMF). Three biological replicates were performed per time point and 3 field of views per replicate were randomly selected for z-stack 3D imaging. The biomass, average and thickness was calculated using the COMSTAT2 (v 2.1) ImageJ (v 2.8.0) plugin (47).

### Protein preparation and LC-MS/MS

Ten micrograms of protein extract from biofilm and planktonic cells were reduced with dithiothreitol, alkylated with iodoacetamide and then digested with trypsin as described in Luu *et al*. (42). The peptides were analysed on the LTQ-Orbitrap Velos mass spectrometer (Thermo Fisher Scientific) at the UNSW Bioanalytical Mass Spectrometry Facility (BMSF) with the settings described in Luu *et al*. (42). The output spectra were matched against a custom *B. pertussis* database on the MaxQuant (v2.0.3.1) proteomics software with the following parameters: digestion mode – specific, enzyme – Trypsin/P, max missed cleavages – 1, Label free quantification – LFQ, Protein identification false discovery rate – 0.01 and min peptides per protein – 2. All other parameters were set as the recommended default values. Student’s *t*-test was calculated and a false discovery rate (FDR) *q*-value multiple test correction was performed using the Storey-Tibshirani method on R (v4.1.1) (48). Proteins were considered upregulated if the fold change (FC) was > 1.2, *q* < 0.05 and downregulated if FC < 0.8, *q* < 0.05 based on previous studies (40). Functional categories were assigned to proteins based on Bart *et al*. (49). The mass spectrometry proteomics data have been deposited to the ProteomeXchange Consortium via the PRIDE (50) partner repository with the dataset identifier PXD033664 and 10.6019/PXD033664.

### Integrative Metabolic Analysis Tool (iMAT) model generation

The proteomic expression data was used to generate context specific metabolic models for the planktonic and biofilm cells. The iMAT (35) method, available in the COBRA Toolbox (v3.0) (51), was used. Processing was done in MATLAB (R2020a) using the IBM CPLEX optimiser (v12.10.0). The iMAT algorithm extracts a simplified model based on the trade-off between high expression and low expression reactions. The comprehensive, manually curated *B. pertussis* metabolic model (iBP1870) generated by Branco dos Santos *et al*. (31) was used as the base model. Subsystems from the *Escherichia coli* genome scale metabolic model (iAF1260) were assigned to each of the reactions in the *B. pertussis* model (52). The *E. coli* (iAF1260) model was used as a template for the original *B. pertussis* model (iBP1870) and therefore, most of the subsystems are transferrable. The original *B. pertussis* iBP1870 model had the tricarboxylic acid (TCA) cycle partially dysfunctional (no flux from oxaloacetate to ⍰-ketoglutarate), as this model was based on previous studies which stated *B. pertussis* has an incomplete TCA cycle (41). However, recent studies have shown that the TCA cycle is fully functional and therefore the disabled pathways were activated before generating the iMAT models (53). After refining the *B. pertussis* model, the biofilm and planktonic proteins were assigned to the gene-protein-reaction (GPR) associations listed in the *B. pertussis* model (iBP1870).

Label free quantification (LFQ) intensity values designated by the MaxQuant software were used to estimate relative protein abundance. To increase confidence in the imported data, only proteins identified in all six biological replicates per condition were incorporated. Proteins were designated as uniquely identified if found in all 6 biological replicates in one condition and none in the other. The expression data was defined as highly (1), lowly (−1) or moderately (0) expressed based on the threshold of mean expression ± 0.5 x STD of the proteins included in the model (35). After generating the planktonic and biofilm context specific iMAT models, the reactions in the biofilm and planktonic models were compared (Figure 1).

**Figure 1.**
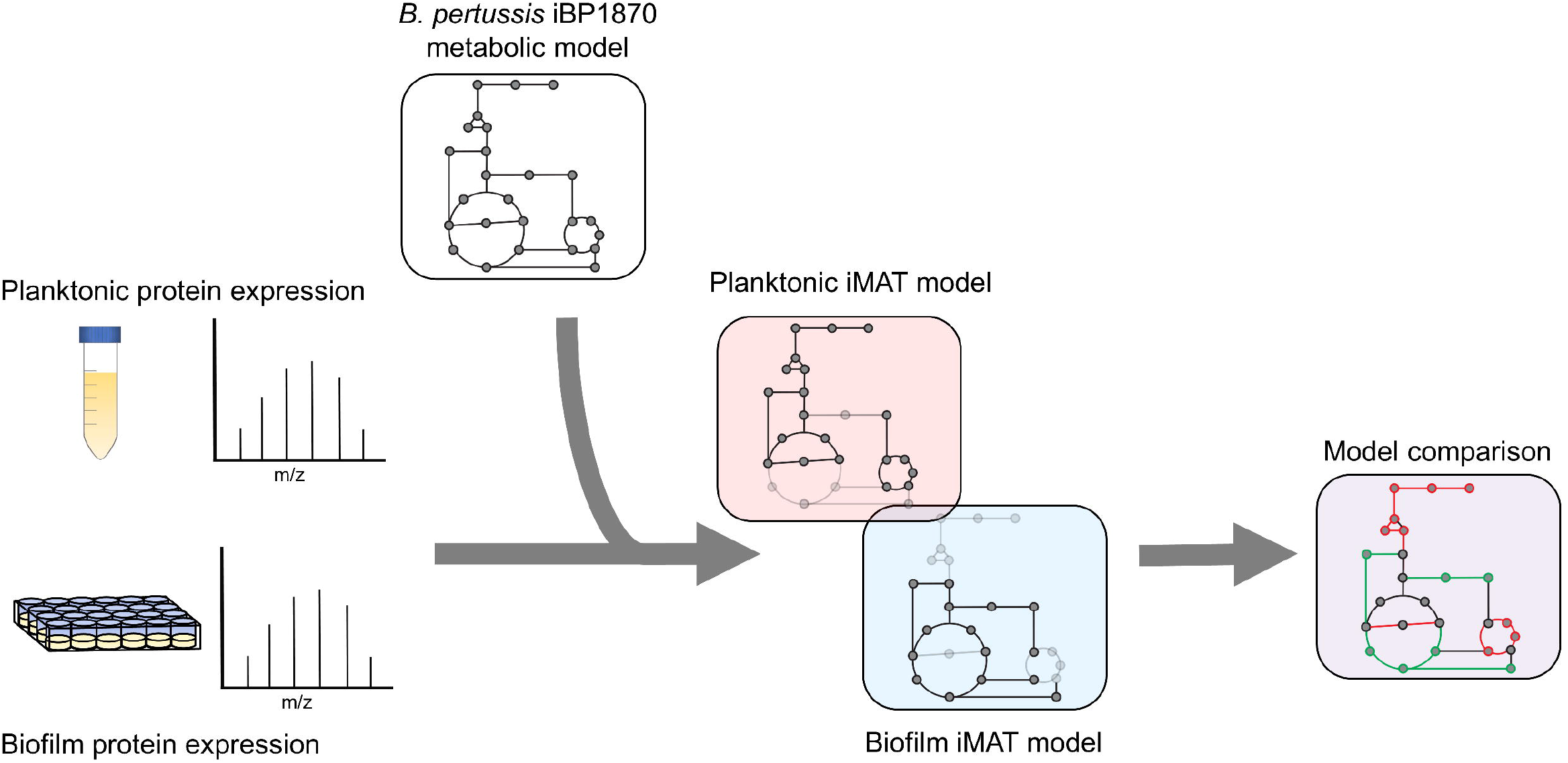
Experimental design for iMAT model generation. Proteins were extracted from planktonic and biofilm *B. pertussis* cells. The expression values were then used to generate context specific metabolic models based on the extensively curated iBP1870 *B. pertussis* model (322). Flux balance analysis was run on the models and the altered reactions compared between the planktonic and biofilm models.

Additional models were generated with proteomic expression data from a separate study. To our knowledge, there are 5 studies that have compared protein expression of *B. pertussis* biofilm cells with their planktonic counterpart (14–18). Of these, only one study by de Gouw *et al*. (14) has publicly available global proteomic expression data of planktonic and biofilm cells and thus was chosen for the production of iMAT models. The proteomic study by de Gouw *et al*. (14) compared *B. pertussis* biofilm and planktonic (mid-log and stationery) cytosolic and membrane proteins. The averaged expression values of the cytosolic and membrane fractionated proteins were combined, and mid-log planktonic expression values were used to generate the models. All models have been deposited to the BioModels database (54) in SBML L3V1 format under the model identifier MODEL2205270001. Differences between the growth conditions are listed in Supplementary Table S1.

### Flux Analysis

A flux variability analysis (FVA) (55) using the COBRA Toolbox (v3.0) (51) was performed on the models which provides minimum and maximum permissible flux bounds for each reaction. Flux is recorded as a millimolar concentration of metabolite per gram of dry cell weight per hour (mmol · g_DCW_^-1^ · h^-1^). To compare the similarity in flux bounds between the biofilm and planktonic models, the Jaccard index function within the COBRA Toolbox (v3.0) (51) was used. This process assigns a similarity index (1 = most similar) by comparing the minimum and maximum flux bounds for the common reactions between the two models for each individual reaction. Many of the reactions have a forward and/or reverse directionality. This is represented in the model as a positive (forward) and negative (reverse) flux value. The FVA values are helpful in determining the predicted directionality of the reaction in the model as flux bounds would often be limited to negative or positive values.

Each model contains a complete set of reactions, however, not all reactions are essential to the model. If a reaction that is essential to the model is missing, the FVA function in the COBRA Toolbox (v3.0) (51) states that the model is infeasible. To identify the most important reactions in each model, each reaction was individually removed in turn and an FVA calculation was attempted to determine whether the model would still be feasible without the reaction.

A flux balance analysis (FBA) was performed using COBRA Toolbox (v3.0) (51) to identify changes in flux in reactions shared by the planktonic and biofilm models. The FBA provides a single value for each reaction based on a predefined metabolic goal so that the models may be compared. The FBA method has been extensively developed to predict metabolic fluxes within metabolic models (24). Typically, the objective function (*c*) of the linear optimisation equation is set as an artificial biomass reaction as the metabolic goal of the organism. As generating biomass may not be the metabolic objective of biofilm cells, an alternate method for defining the objective function was used in this study. The objective function was defined based on proteomic expression data as described in Montezano *et al*. (56). For all proteins that were identified with GPR associations in the model, the intensities were normalised by the maximum intensity value for each condition (planktonic and biofilm) and these values were input as *c*. This leads the FBA to push flux towards the reactions which have higher protein expression and considerably shrinks the solution space to increase prediction accuracy (56).

## Results

### Key proteomic changes identified between biofilm and planktonic cells

To identify changes that occur in biofilm conditions, label free quantification mass spectrometry (LFQ-MS) was performed on biofilm and planktonic cells. Analysis was performed on L1423, a clinical isolate representative of the current circulating *B. pertussis* strains. Confocal microscopy confirmed biofilm formation and mature structure at 96 h (Supplementary Figure S1). Furthermore, there was a polysaccharide biosynthesis protein, WbpO (FC = 2.98, *q* = 1.17E-5), a phosphoglucomutase enzyme, Pgm (FC = 1.23, *q* = 0.041) and an outer membrane porin protein, BP0840 (FC = 9.45, *q* = 1.05E-5) that were seen to be upregulated in biofilm cells (Figure 2B). These proteins have been previously linked with *B. pertussis* biofilm reinforcing the biofilm phenotype achieved in this study (18). There were 948 proteins identified in total (Supplementary Table S2), of which, 571 were proteins identified in all 6 biological replicates (Figure 2A). There were 478 proteins with significantly differential expression (*q* < 0.05) between the two conditions. In biofilm cells, there were 242 proteins downregulated and 236 proteins upregulated (Figure 2B).

**Figure 2.**
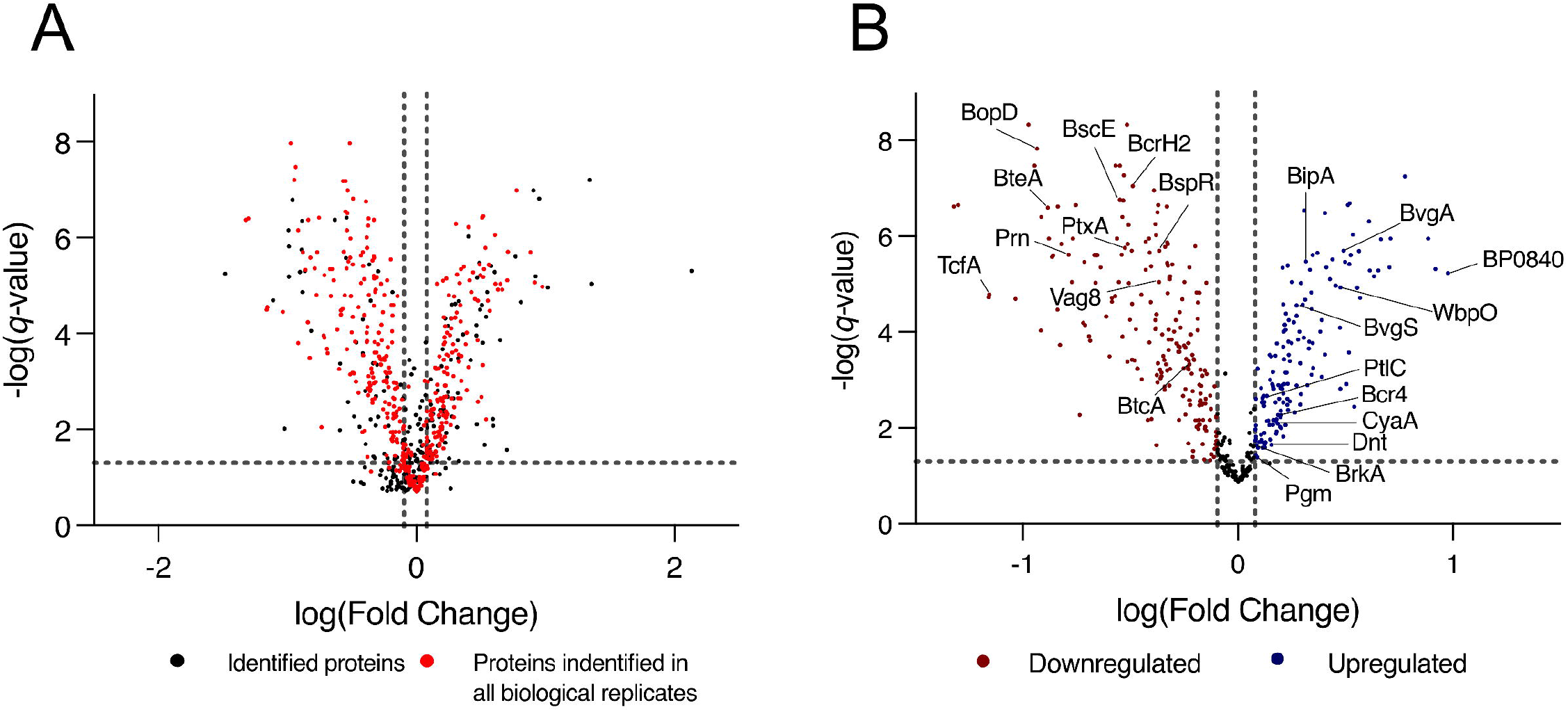
Volcano plot of total protein expression changes between biofilm and planktonic cells. **A** Expression profile of all proteins identified through the LC-MS/MS analysis of planktonic and biofilm *B. pertussis* cells. Proteins are plotted on a volcano plot displaying the -log(*q-* value) on the y-axis and log(fold change [biofilm/planktonic]) on the x-axis. The dashed vertical grey lines mark a fold change of 0.8 and 1.2 and the horizontal line marks the threshold of *q*-value = 0.05. Highlighted in red are the proteins that were identified in all 6 biological replicates that were incorporated into the iMAT models. **B** Expression profile of the subset of proteins incorporated into the iMAT model. Red markers are proteins that were designated significantly differentially downregulated, and the blue markers are proteins that were significantly upregulated in biofilm cells. The dashed vertical grey lines mark a fold change of 0.8 and 1.2 and the horizontal line marks the threshold of *q*-value = 0.05. Proteins of interest are labelled.

There were many proteins related to virulence with altered expression identified in this study (Table 1). BipA, an outer membrane protein associated with biofilm formation was upregulated (14). Adenylate cyclase toxin (CyaA) and dermonecrotic toxin (Dnt) were also upregulated in biofilm (Figure 2B). There were 10 proteins from the Type III secretion system (T3SS) that were downregulated in biofilm cells or uniquely identified in planktonic cells. One protein (Bcr4) involved in the T3SS was upregulated in biofilm cells (57). Tracheal colonisation factor (TcfA) and virulence associated gene 8 (Vag8) were downregulated in biofilm cells while *Bordetella* resistance to killing (BrkA) protein was upregulated (Figure 2B). Of the ACV antigens, there was no change in expression for filamentous haemagglutinin (FhaB) but the filamentous haemagglutinin outer membrane transporter protein (FhaC) was uniquely identified in biofilm cells. The membrane bound pertussis toxin subunit 4 (PtxD) was also uniquely identified in biofilm cells, however, pertussis toxin subunit 1 (PtxA) was downregulated in biofilms. Furthermore, fimbriae protein (Fim2) was downregulated in biofilm. Finally, pertactin (Prn) was strongly downregulated in biofilm (FC = 0.16, *q* = 2.42E-6). These changes demonstrate a strongly altered virulence profile in *B. pertussis* biofilm cells compared to planktonic cells.

**Table 1.**
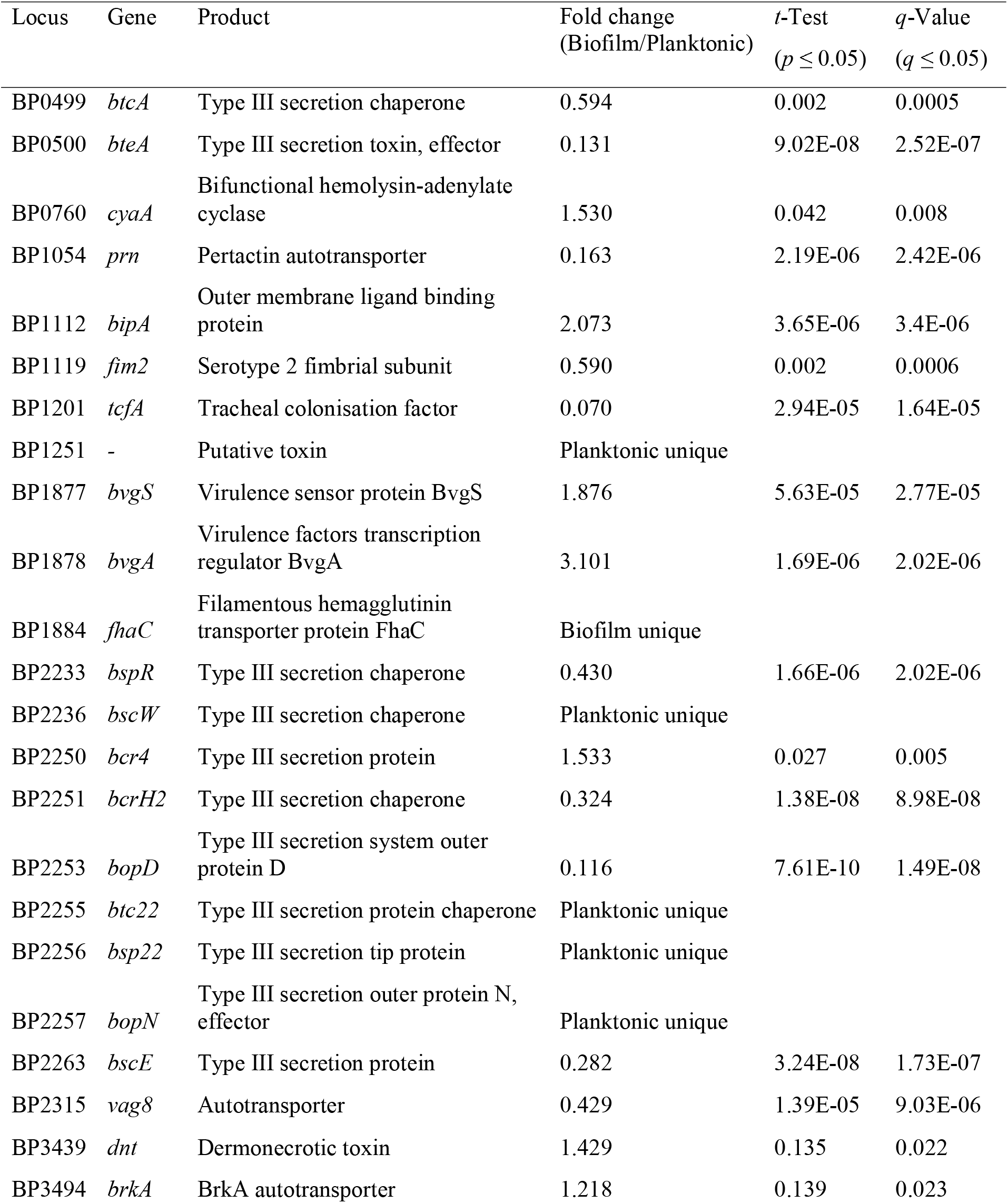
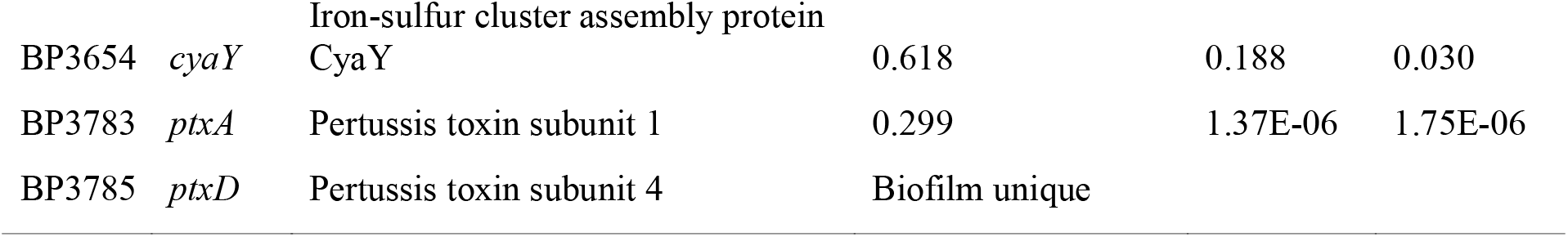
Virulence proteins differentially expressed or uniquely identified between planktonic and biofilm cells

| Locus | Gene | Product | Fold change<br>(Biofilm/Planktonic) | *t*-Test<br>($p \leq 0.05$) | *q*-Value<br>($q \leq 0.05$) |
| --- | --- | --- | --- | --- | --- |
| BP0499 | *btcA* | Type III secretion chaperone | 0.594 | 0.002 | 0.0005 |
| BP0500 | *bteA* | Type III secretion toxin, effector | 0.131 | 9.02E-08 | 2.52E-07 |
| BP0760 | *cyaA* | Bifunctional hemolysin-adenylate cyclase | 1.530 | 0.042 | 0.008 |
| BP1054 | *prn* | Pertactin autotransporter | 0.163 | 2.19E-06 | 2.42E-06 |
| BP1112 | *bipA* | Outer membrane ligand binding protein | 2.073 | 3.65E-06 | 3.4E-06 |
| BP1119 | *fim2* | Serotype 2 fimbrial subunit | 0.590 | 0.002 | 0.0006 |
| BP1201 | *tcfA* | Tracheal colonisation factor | 0.070 | 2.94E-05 | 1.64E-05 |
| BP1251 | - | Putative toxin | Planktonic unique |  |  |
| BP1877 | *bvgS* | Virulence sensor protein BvgS | 1.876 | 5.63E-05 | 2.77E-05 |
| BP1878 | *bvgA* | Virulence factors transcription regulator BvgA | 3.101 | 1.69E-06 | 2.02E-06 |
| BP1884 | *fhaC* | Filamentous hemagglutinin transporter protein FhaC | Biofilm unique |  |  |
| BP2233 | *bspR* | Type III secretion chaperone | 0.430 | 1.66E-06 | 2.02E-06 |
| BP2236 | *bscW* | Type III secretion chaperone | Planktonic unique |  |  |
| BP2250 | *bcr4* | Type III secretion protein | 1.533 | 0.027 | 0.005 |
| BP2251 | *bcrH2* | Type III secretion chaperone | 0.324 | 1.38E-08 | 8.98E-08 |
| BP2253 | *bopD* | Type III secretion system outer protein D | 0.116 | 7.61E-10 | 1.49E-08 |
| BP2255 | *btc22* | Type III secretion protein chaperone | Planktonic unique |  |  |
| BP2256 | *bsp22* | Type III secretion tip protein | Planktonic unique |  |  |
| BP2257 | *bopN* | Type III secretion outer protein N, effector | Planktonic unique |  |  |
| BP2263 | *bscE* | Type III secretion protein | 0.282 | 3.24E-08 | 1.73E-07 |
| BP2315 | *vag8* | Autotransporter | 0.429 | 1.39E-05 | 9.03E-06 |
| BP3439 | *dnt* | Dermonecrotic toxin | 1.429 | 0.135 | 0.022 |
| BP3494 | *brkA* | BrkA autotransporter | 1.218 | 0.139 | 0.023 |
| BP3654 | *cyaY* | Iron-sulfur cluster assembly protein<br>CyaY | 0.618 | 0.188 | 0.030 |
| BP3783 | *ptxA* | Pertussis toxin subunit 1 | 0.299 | 1.37E-06 | 1.75E-06 |
| BP3785 | *ptxD* | Pertussis toxin subunit 4 | Biofilm unique |  |  |

When the proteins were grouped into functional categories based on Bart *et al*. (49), there was a significant increase in proteins in the categories of transport/binding and miscellaneous proteins under biofilm conditions. There was also a downregulation of the functional groups: ribosome constituents and cell processes (Figure 3).

**Figure 3.**
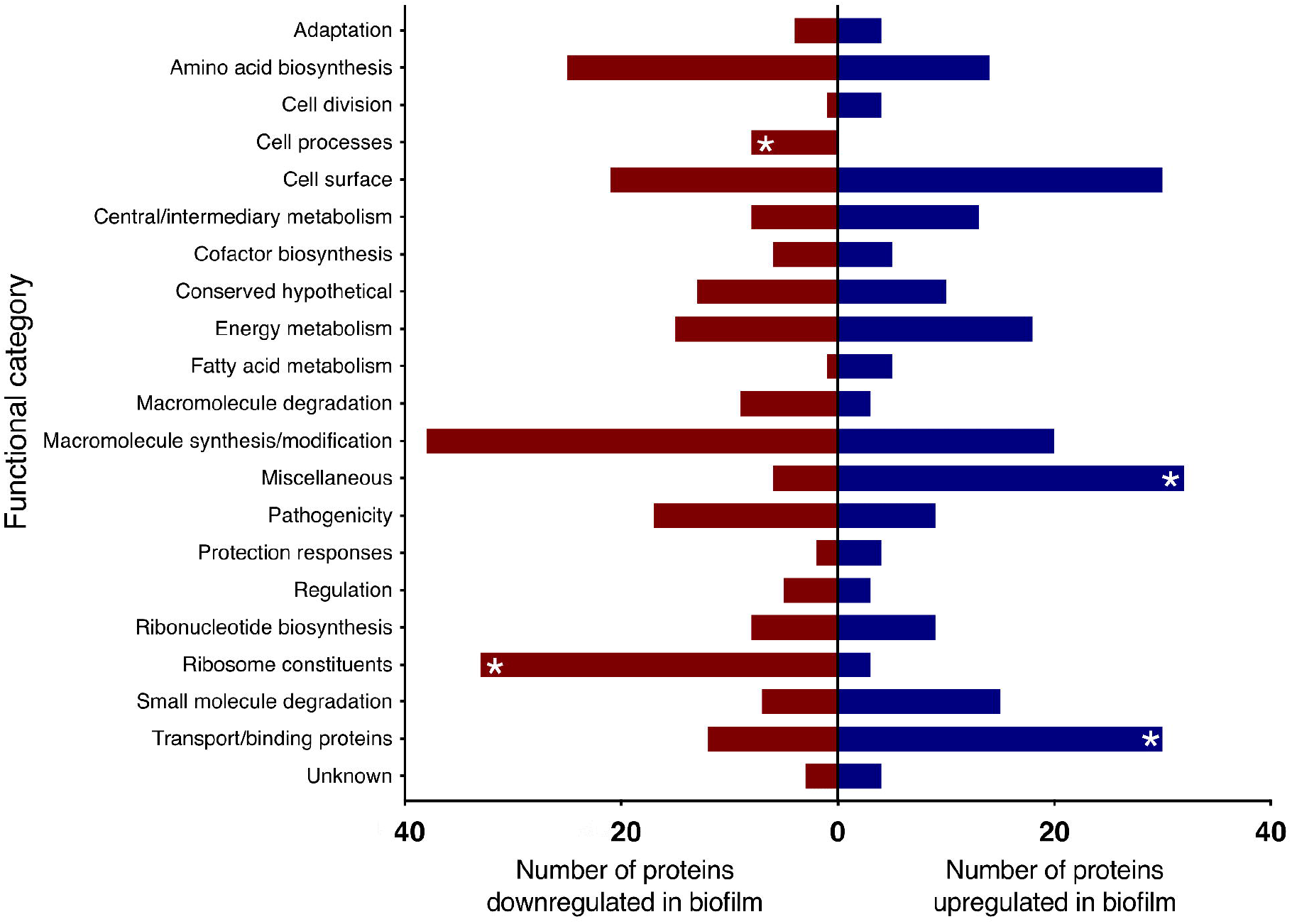
Proteins up and downregulated in *B. pertussis* biofilm cells compared to planktonic cells identified using LC-MS/MS. Proteins significantly up and downregulated in biofilm cells were categorised in functional categories based on Bart *et al*. (510). Red and blue bars represent the total number of proteins within the functional category significantly up or downregulated, respectively. Asterisk (*) denotes functional categories significantly up or downregulated based on Fisher’s exact test with Benjamini-Hochberg multiple test correction (adjusted *p* < 0.05).

### iMAT model generation

For an in-depth analysis of the metabolic changes between biofilm and planktonic cells, iMAT metabolic models were generated based on protein expression from both conditions. Of the 571 proteins identified using mass spectrometry, 228 were annotated with GPR associations in the *B. pertussis* (iBP1870) model (31). These are proteins which have assigned reactions in the base model. The overlap between the identified proteins and the reactions in the model constitutes 29.61% of the total GPR associations listed in the base model. Context specific models were created using the iMAT algorithm implemented in the COBRA Toolbox (51). The biofilm iMAT model consisted of 198 metabolites, 206 reactions and 188 genes. The planktonic iMAT model had 213 metabolites, 219 reactions and 195 genes (Table 2). To identify changes between the planktonic and biofilm iMAT models, the number of unique and common reactions between the two models were compared. There were 168 reactions that were common between the two models, 51 reactions that were unique to the planktonic model and 38 that were unique to the biofilm model (Figure 4A). The reactions were grouped into subsystems and total number of reactions in each group were similar between the two models (Figure 4B). It is notable however that there was a high proportion of reactions that were unique to the individual models. Major pathways are summarised and represented in Figure 5. Additionally, metabolic pathways for the tricarboxylic acid (TCA) cycle, arginine metabolism, aspartate metabolism and glycerophospholipid metabolism pathways are highlighted in Figures 6, 7 and 8.

**Figure 4.**
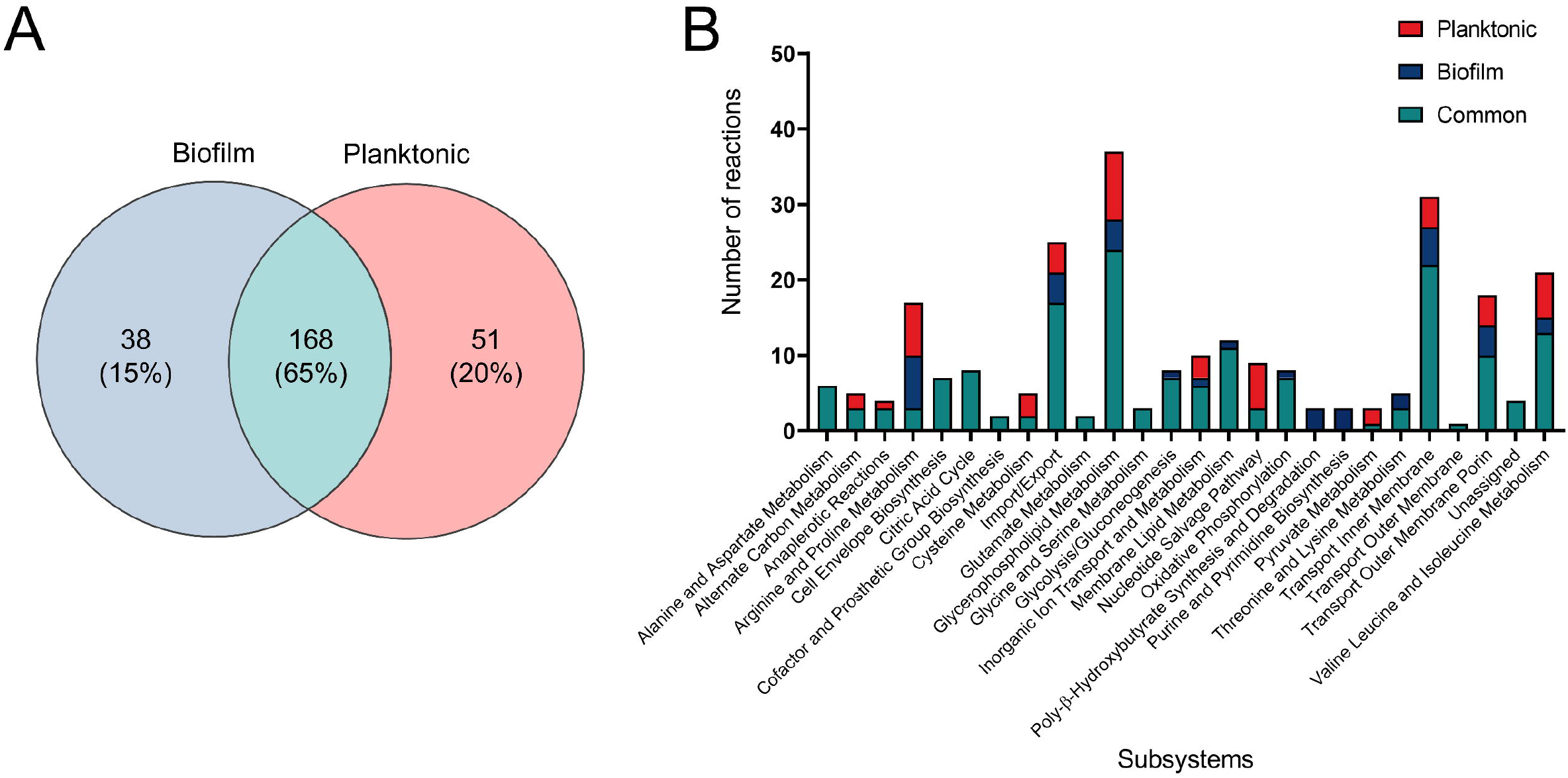
Comparison of reactions of *B. pertussis* planktonic and biofilm iMAT models. **A** Venn diagram highlighting common and unique reactions between planktonic and biofilm iMAT models generated by incorporating proteomic expression data. **B** Planktonic and biofilm model reactions grouped into subsystems based on the *Escherichia coli* iAF1260 metabolic model. Unique reactions identified in one model but not the other are highlighted in different colours on the graph.

**Figure 5.**
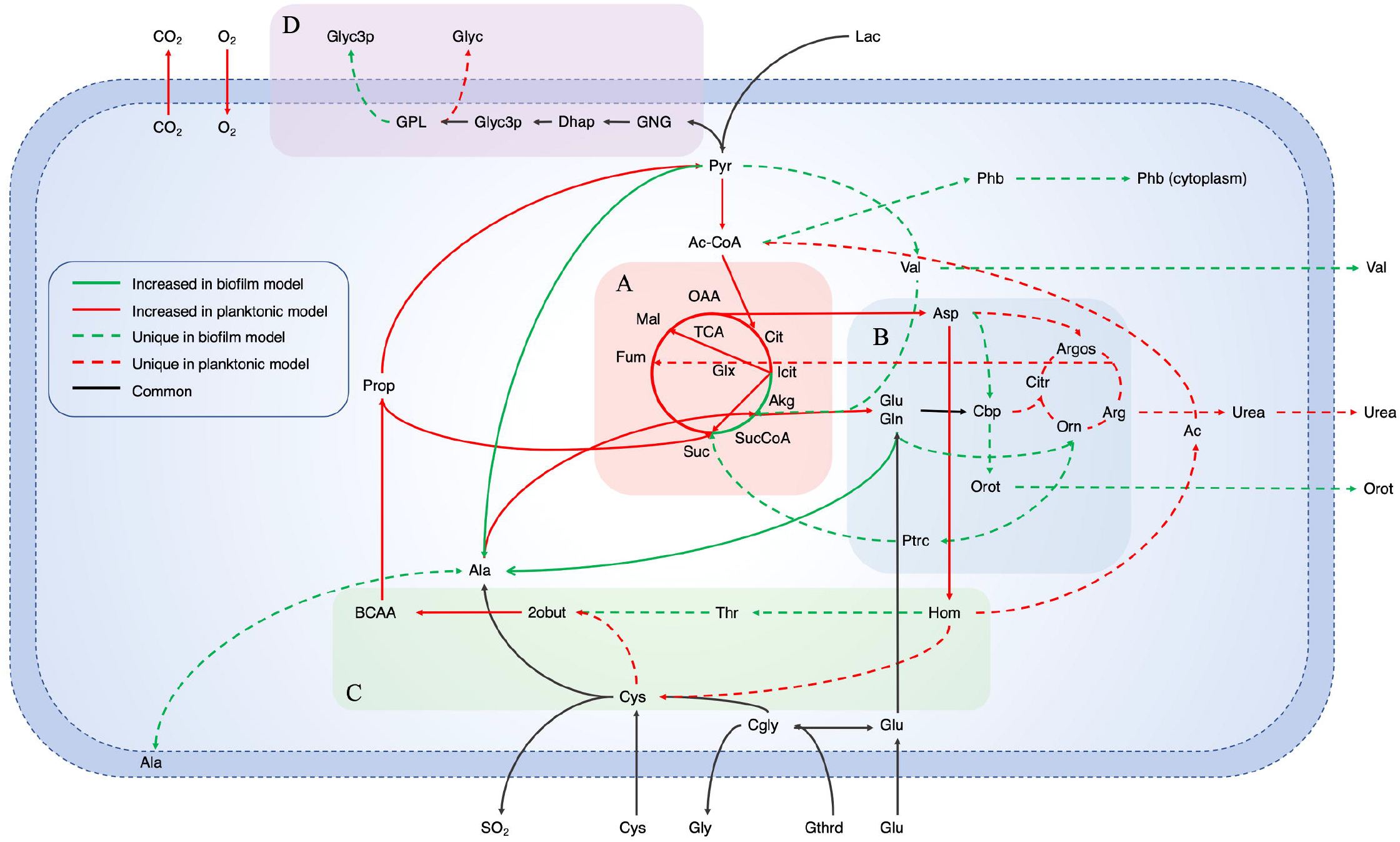
Summarised pathways with major changes between planktonic and biofilm iMAT models. Major changed core metabolic reactions from a comparison of iMAT models generated from biofilm and planktonic *B. pertussis* protein expression data. The model includes the reactions that were unique in either model as well as reactions with altered flux. Red lines indicate reactions that had decreased flux in biofilm cells while green lines indicate reactions with increased flux. The dashed lines indicate unique reactions to each model while the black lines are common reaction with the same flux. Highlighted sections are **A** the Tricarboxylic acid cycle, **B** Arginine metabolism, **C** Aspartate metabolism and **D** Gluconeogenesis and glycerophospholipid metabolism. Pathways for **A**, **B** and **C** are represented in more detail in Figures **6**, **7** and **8**, respectively. Ac, Acetate; Ac-CoA, Acetyl-CoA; Akg, α-ketoglutarate; Ala, Alanine; Arg, Arginine; Argos, Argininosuccinate; Asp, Aspartate; BCAA, Branched chain amino acid degradation; Cbp, Carbamoyl phosphate; Cgly, Cysteinylglycine; Cit, Citrate; Citr, Citruline; CO_2_, Carbon dioxide; Cys, Cysteine; Dhap, Dihydroxyacetone phosphate; Fum, Fumarate; Gln, Glutamine; Glu, Glutamate; Glx, Glyoxylate; Gly, Glycine; Glyc3p, Glycerol 3-phosphate; Glyc, Glycerol; GNG, Gluconeogenesis; GPL, Glycerophospholipid metabolism; Gthrd, Glutathione; Hom, Homoserine; Icit, Isocitrate; Lac, Lactate; Mal, Malate; O_2_, Oxygen; OAA, Oxaloacetate; Orn, Ornithine; Orot, Orotate; Phb, Polyhydroxybutyrate; Prop, Propanoate metabolism; Ptrc, Putrescine; Pyr, Pyruvate; SO_2_, Sulfur dioxide; Suc, Succinate; SucCoA, Succinyl-CoA; TCA, Tricarboxcylic acid cycle; Thr, Threonine; Val, Valine; 2obut, 2-oxobutanoate.

**Figure 6.**
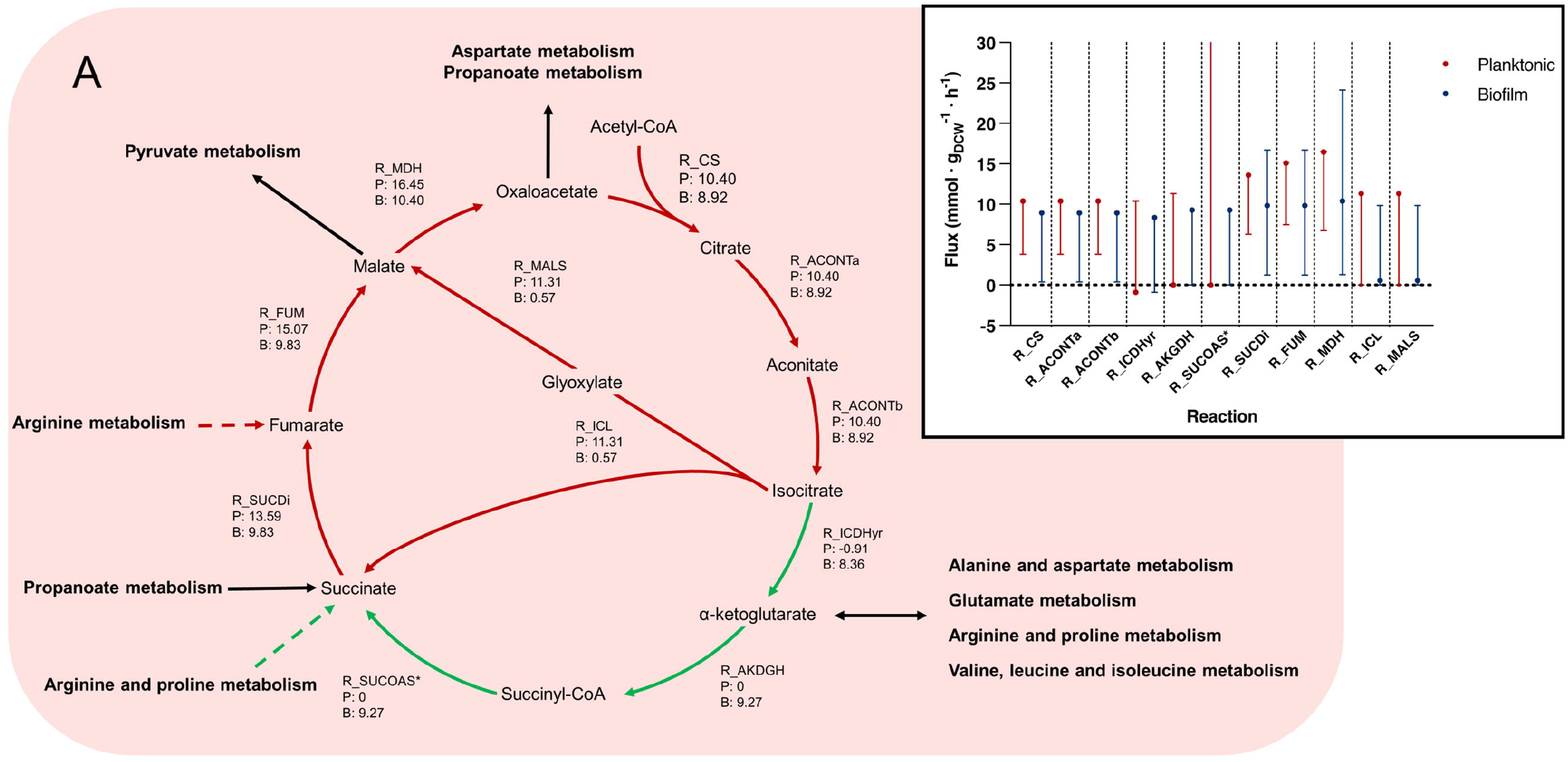
Tricarboxylic acid cycle pathways and flux bounds between planktonic and biofilm iMAT models. This figure relates to Figure 5A. Reactions within the tricarboxylic acid (TCA) cycle between planktonic and biofilm *B. pertussis* iMAT metabolic models generated from proteomic expression data. Reaction names and flux values from the flux balance analysis (FBA) are given for each reaction within the TCA. Each arrow is indicates a reaction and flux values are labelled as (P) representing planktonic model flux and (B) representing biofilm model flux. All flux values are mmol · gDCW ^1^ · h^-1^. Green arrows represent reactions that are upregulated in biofilm cells while red arrows are downregulated reactions. Green dashed arrows are unique reactions to the biofilm model while red dashed arrows are reactions unique to the planktonic model. Black arrows are common reactions. General metabolic pathways are in bold. Inset Range for flux variance analysis and FBA values for the tricarboxylic acid cycle between planktonic and biofilm cells. Flux ranges (min to max) are represented as lines and the FBA value is annotated with a point. Planktonic values are indicated in red and biofilm values in blue. *R_SUCOAS is a reaction that flows from succinate to succinyl-CoA. The reaction runs in reverse in the typical TCA cycle and so these values are negative. For simplicity, these reactions have been annotated as absolute values in the figure. Furthermore, the flux bounds for the planktonic models are at the max value and the max has been omitted from the graph. R_CS, type II citrate synthase; R_ACONTa, citrate hydrolase; R_ACONTb, aconitate hydratase; R_ICDHyr, isocitrate dehydrogenase; R_AKGDH, α-ketoglutarate dehydrogenase; R_SUCOAS, succinyl-CoA synthetase; R_SUCDi, succinate dehydrogenase; R_FUM, fumarate hydratase; R_MDH, malate dehydrogenase; R_ICL, isocitrate lyase; R_MALS, malate synthase.

**Figure 7.**
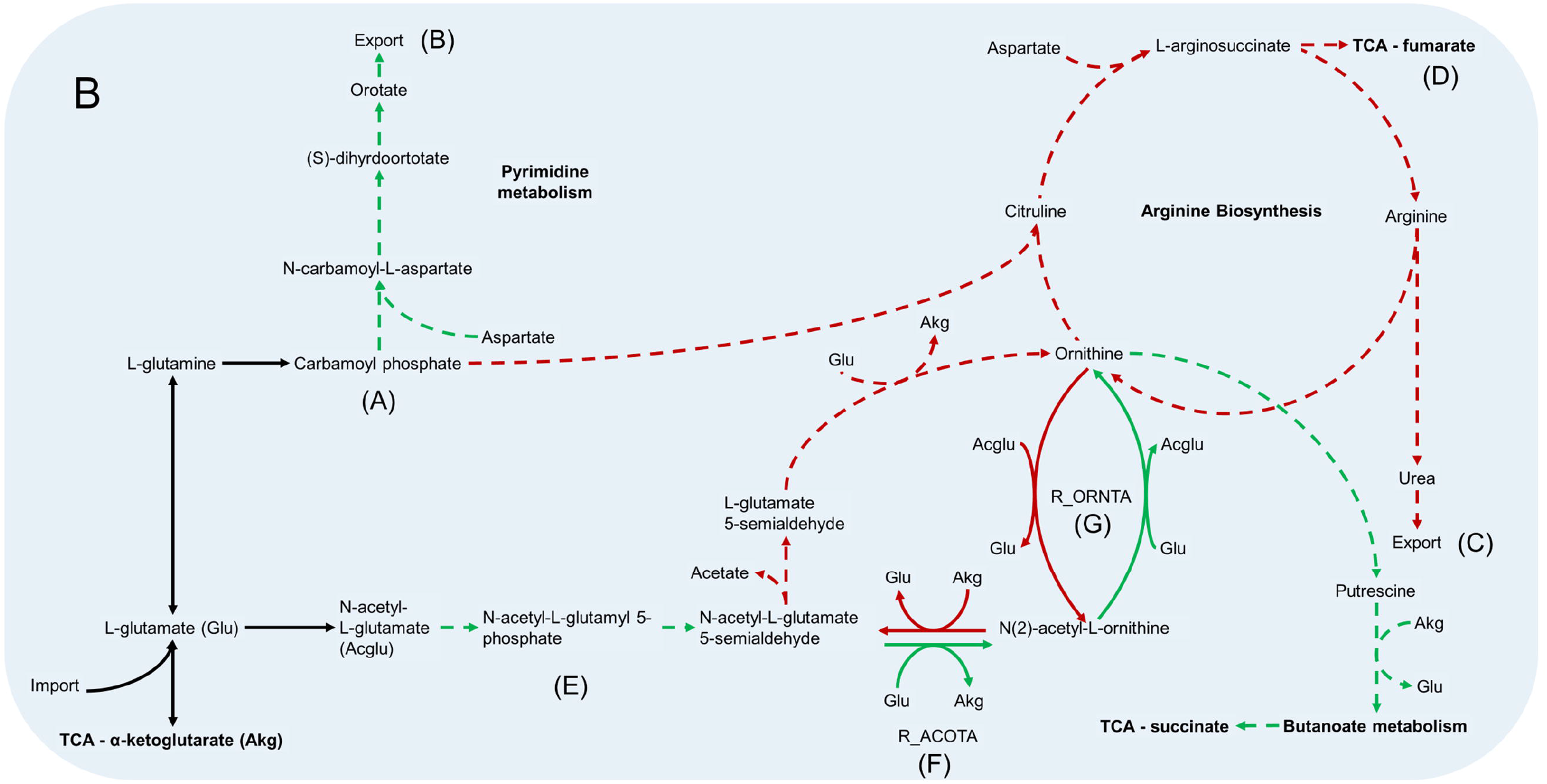
Arginine metabolism pathways in *B. pertussis* biofilm and planktonic iMAT models. This figure relates to Figure 5B. Models were generated using proteomic expression data. Green arrows represent reactions that are upregulated in biofilm cells while red arrows are downregulated reactions. Green dashed arrows are unique reactions to the biofilm model while red dashed arrows are reactions unique to the planktonic model. Black arrows are common reactions. General metabolic pathways are in bold. 0-ketoglutarate (Akg) and glutamate (Glu) are utilised in many of the reactions and are therefore included multiple times in abbreviated form. N-acetyl-L-glutamate is also included as the abbreviation Acglu for the reaction R_ORNTA. Key sections of the pathways have also been annotated. **A** Both models generate carbamoyl phosphate through the same reactions. **B** The biofilm model has unique reactions to synthesise orotate from carbamoyl phosphate and export it out the cell. **C** The planktonic model completed the arginine biosynthesis pathway and exports urea out of the cell. **D** In addition to the urea, the arginine biosynthesis pathway leads to the production of fumarate which is fed back into the TCA. **E** The pathways from N-acetyl-L-glutamate to N-acetyl-L-glutamate 5 semialdehyde are unique to the biofilm model. **F** The reaction, acetylornithine transaminase reaction (R_ACOTA), runs in opposite directions for planktonic and biofilm models. **G** The reaction, glutamate N-acetyl transferase (R_ORNTA), also runs in opposite directions for biofilm and planktonic models.

**Figure 8.**
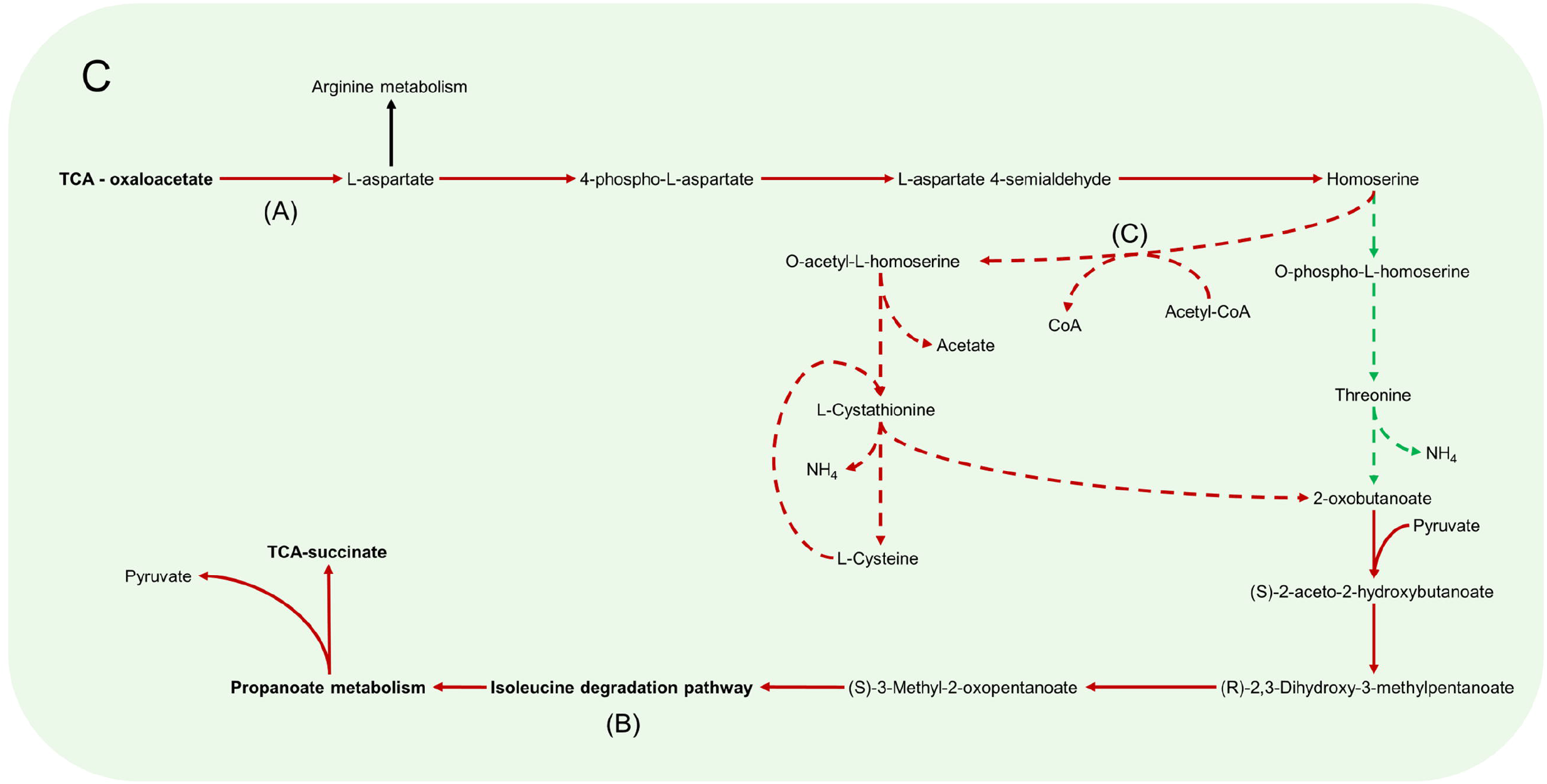
Aspartate metabolism pathways in *B. pertussis* biofilm and planktonic iMAT models. This figure relates to Figure 5C. Models were generated using proteomic expression data. Red arrows represent downregulated reactions in the biofilm model. Green dashed arrows are unique reactions to the biofilm model while red dashed arrows are reactions unique to the planktonic model. Black arrows are common reactions. General metabolic pathways are in bold. Key sections of the pathways have also been annotated. **A** There is a decreased flux from oxaloacetate to L-aspartate in the biofilm model. **B** In both models, 2-oxobutanoate is eventually degraded through the branched chain amino acid degradation pathway specifically, the isoleucine pathway. **C** While biofilm cells convert homoserine through to O-phospho-L-homoserine, planktonic cells convert homoserine to O-acetyl-L-homoserine by utilising acetyl-CoA.

**Table 2.** Number of metabolites, reactions and genes in the iMAT context specific models for *B. pertussis* biofilm and planktonic cells.

|  | Biofilm | Planktonic |
| --- | --- | --- |
| High expression (+1) | 36 | 36 |
| Moderate expression (0) | 117 | 86 |
| Low expression (-1) | 65 | 78 |
| Total proteins incorporated | 218 | 200 |
| **iMAT models** |  |  |
| Metabolites | 198 | 213 |
| Reactions | 206 | 219 |
| Genes | 188 | 195 |

### *B. pertussis* biofilm cells complete the tricarboxylic acid cycle

An FBA was performed to reveal changes in flux between reactions that were common between the planktonic and biofilm models. A major difference was seen in the TCA cycle. The biofilm cells pushed flux to complete the TCA cycle while the planktonic cells pushed flux towards the glyoxylate shunt (Figure 5A). While there was slightly higher levels of flux from oxaloacetate through to isocitrate in the planktonic model (planktonic: 10.40 mmol · g_DCW_^-1^ · h^-1^, biofilm: 8.92 mmol · g_DCW_^-1^ · h^-1^), the planktonic cells pushed all flux (11.31 mmol · g_DCW_^-1^ · h^-1^) into the glyoxylate shunt to convert isocitrate to glyoxylate and succinate while comparatively, only a small flux (0.57 mmol · g_DCW_^-1^ · h^-1^) was pushed for the same reactions for biofilm cells (Figure 6). Instead, the flux moved towards ⍰-ketoglutarate (AKG) to succinyl-CoA and succinate to complete the TCA cycle (Figure 6). The planktonic cells comparatively had no flux through these reactions. Despite these changes, there was an increased level of flux from succinate to fumarate and malate in the planktonic model which was fed from the glyoxylate shunt, branched chain amino acid degradation and the arginine biosynthesis cycle. Finally, there was also increased conversion of pyruvate to acetyl-CoA in the planktonic model which was then fed back into the TCA cycle as citrate. Most of the acetyl-CoA synthesis/utilisation pathways were either unique to the planktonic model or downregulated in the biofilm model. However, the biofilm model had unique reactions to convert acetyl-CoA to polyhydroxybutyrate (PHB) (Figure 5).

### Cardiolipin synthesis active in planktonic cells

There were many reactions included in both models that were grouped into the glycerophospholipid metabolism subsystem. Flux moves through the gluconeogenesis pathway in both the models from pyruvate through to dihydroxyacetone phosphate and then glycerol 3-phosphate (Figure 5D). The pathways for most of the glycerophospholipids and fatty acid synthesis were the same between the two models. However, the planktonic model had pathways producing cardiolipin (tetradodecanoyl, n-C12:0) while the biofilm model lacked these pathways. The production of cardiolipin is through the pathway of conversion of 1,2-didodecanoyl-sn-glycero-3-cytidine 5’-diphosphate (CDP-DAG) to phosphatidylglycerophosphate (didodecanoyl, n-C12:0) and therefore there is a higher level of flux from the cytidine monophosphate (CMP) to cytidine triphosphate (CTP) in the planktonic model. The models suggest that there would be a higher level of 1-dodecanoyl-*sn*-glycerol 3-phosphate, 1-hexadec-9-enoyl-*sn*-glycerol 3-phosphate and glycerol 3-phosphate in the periplasm of the biofilm cells. In the planktonic cells, there was a higher level of cardiolipin, phosphatidylglycerol (didodecanoyl, n-C12:0), 2-dodecanoyl-*sn*-glycerol 3-phosphate and 2-hexadec-9-enoyl-*sn*-glycerol 3-phosphate. The biofilm model uniquely exported glycerol 3-phosphate into the extracellular space through these pathways while planktonic model uniquely exported glycerol into the extracellular space (Figure 5D).

### Arginine biosynthesis decreased in biofilm cells

The metabolic models also revealed many altered amino acid metabolism pathways. There was an equal number of reactions grouped into the arginine and proline metabolism subsystem between the two models. However, the processes within the pathways were varied as seen by the number of unique reactions within the subsystem (Figure 4B). Both models feed into the arginine metabolism pathways through conversion of L-glutamate (GLU). Subsequently, both models have reactions to produce carbamoyl phosphate (CBP) (Figure 7A). While the planktonic model pushed the CBP into the arginine biosynthesis pathway, the biofilm model had reactions to convert CBP through to orotate which was then exported out of the cell (Figure 7B). The reactions surrounding orotate were grouped in the purine and pyrimidine metabolism subsystem unique to the biofilm model (Figure 4B). The planktonic cells complete the arginine biosynthesis cycle, exporting urea out of the cell as a by-product (Figure 7C). Furthermore, in the planktonic model, argininosuccinate is converted to arginine and fumarate which is fed back into the TCA cycle (Figure 5A and Figure 7D). When each of these reactions were deleted in turn and tested for model feasibility, it led to infeasible models for the planktonic model but were still feasible for the biofilm model (Supplementary Table S3).

Both models produced ornithine surrounding the arginine metabolism pathways. The planktonic cells produced ornithine through the arginine biosynthesis pathway while the biofilm cells synthesised ornithine through N-acetyl-L-glutamate to acetyl-ornithine and then to ornithine (Figure 7E). The fate of ornithine also differed greatly between the two models. When the FVA values were compared, the smallest Jaccard index value (lowest similarity) between the models were for the reactions acetylornithine transaminase (R_ACOTA) and glutamate N-acetyl transferase (R_ORNTA). R_ACOTA is a reversible reaction of acetylornithine and AKG to N-acetyl-L-glutamate 5-semialdehyde and GLU. The FVA revealed that the biofilm model only had capabilities to run this reaction in reverse (Figure 7F). The opposite was seen for R_ORNTA, a reversible reaction of acetyl-ornithine and GLU to ornithine and N-acetyl-L-glutamate (Figure 7G). The planktonic model ran this reaction in reverse. Overall, the ornithine in the planktonic model was either utilised to produce acetate or pushed back through the arginine biosynthesis pathway. The biofilm model pushed ornithine through the butanoate metabolism pathway into the TCA cycle as succinate with the production of NADPH and NADH. This process utilised AKG and produced GLU.

### Aspartate metabolism downregulated in biofilm cells

Linked with the arginine metabolism pathways was the amino acid, aspartate (Figure 5C). Aspartate is synthesised in the cells through the aspartate transaminase reaction which converts oxaloacetate and GLU to L-aspartate and AKG. This reaction had decreased flux in the biofilm cells (Figure 8A). Interestingly, when the reaction was deleted in the planktonic model, it led to an infeasible model while the biofilm model remained feasible without the reaction. The flux in the biofilm model pushed from aspartate through to threonine synthesis and CBP metabolism. Threonine is eventually degraded through the branched chain amino acid (BCAA) degradation pathway (Figure 8B). Additionally, aspartate was uniquely converted with CBP to N-carbamoyl-L-aspartate and ultimately into orotate as mentioned above for the biofilm model (Figure 8B). The planktonic model had flux flow from aspartate to homoserine, but rather than following through to threonine, the homoserine was converted to O-acetyl-L-homoserine towards acetate and L-cystathionine (Figure 8C). L-cystathionine was then converted to L-cysteine and 2-oxobutanoate which was fed through the BCAA (specifically the isoleucine pathway) degradation pathway that was upregulated in the planktonic model. The BCAA degradation pathway leads to the production of acetyl-CoA, NADH, NADPH and FADH_2_. There was further production of these molecules through the valine degradation pathway in the planktonic model. These pathways lead flux being moved back into the TCA cycle as succinate and pyruvate through propanoate metabolism. Although the planktonic model moved flux through the valine degradation pathway, the actual production of valine was unique to the biofilm model. The valine was either exported out of the cell or converted to AKG in the biofilm model.

Cysteine was transported into the cell at the same rate between the two models. Both models source cysteine through diffusion, active ABC transport and the conversion of glutathione and L-cysteinylglycine. The biofilm model pushed all flux of cysteine to L-alanine while the planktonic model also has pathways to convert the cysteine through to acetyl-CoA as mentioned above (Figure 8C). The reactions to move L-alanine in and out of the periplasm were unique to the biofilm model. The main source of L-alanine for the biofilm cells was from the conversion of pyruvate and glutamine to L-alanine and AKG. The planktonic model creates L-alanine through the conversion of β-alanine. This reaction also creates malonate-semialdehyde and was strongly downregulated in the biofilm model. The L-alanine is converted to D-alanine and then to GLU and pyruvate in both models.

### Biofilm cells have increased superoxide dismutase activity

Both models converted ubiquinol to ubiquinone through cytochrome ubiquinol oxidase. This reaction was downregulated in biofilm model. However, the biofilm model also had an additional reaction that converted ubiquinol to ubiquinone with the by-product of superoxide anions. The superoxide was converted to hydrogen peroxide and O_2_ by superoxide dismutase. The hydrogen peroxide was converted to H_2_O by thioredoxin. The other reactions involving ubiquinone were downregulated in biofilm cells including succinate dehydrogenase and NADH dehydrogenase. However, ubiquinone was converted to ubiquinol through the conversion S-dihydroorotate to orotate in a reaction unique to the biofilm cells.

### Comparison with other metabolic models

We further used proteomics data from a previous *B. pertussis* biofilm study by de Gouw *et al*. (14) to create iMAT models and compare the reactions. The study by de Gouw *et al*. (14) grew *B. pertussis* biofilms on flat polypropylene beads in a glass column reactor for 72 h with THIJS media refreshed every 24 h. The planktonic cells were extracted at both mid-exponential phase at 17 h and stationary phase at 40 h. That study identified 729 – 825 proteins from the three different conditions with an overlap of 645 proteins (14). While the biofilm iMAT model generated using that proteomic data had a higher number of reactions, metabolites and genes to the planktonic iMAT model generated from that same data, the overall values were comparable to the models of this study (Supplementary Table S1).

In line with our study, the de Gouw *et al*. (14) biofilm model completed the full TCA cycle while the planktonic model pushed flux through the glyoxylate shunt. Further similarities were identified between the biofilm models in NADH dehydrogenase, aspartate transaminase (Figure 8A) and CO_2_ and O_2_ exchange. All biofilm models had decreased flux in these reactions compared to their planktonic counterpart. The NADH dehydrogenase reaction produces NAD+ from NADH with the conversion of ubiquinone to ubiquinol. The aspartate transaminase reaction (R_ASPTA - reversed) had decreased flux for all biofilm models. This reaction converted oxaloacetate and GLU to AKG and L-aspartate. The reaction from aspartate to CBP and subsequent reactions to orotate were unique to the biofilm models (Figure 7B and Supplementary Table S3).

## Discussion

Recent studies have shown that biofilms are an important aspect of *B. pertussis* pathogenesis (17, 46, 58). Biofilm cells are more resilient against antimicrobials and environmental stresses (13, 15). While previous studies of *B. pertussis* biofilms have made important discoveries related to growth and virulence, there has been less research focused on biofilm metabolism. Biofilms are readily formed by *B. pertussis* and has been defined as an integral part of its pathogenesis (7). Investigating the metabolic changes that occur within a biofilm community may help understand the pathogenesis of *B. pertussis*. Therefore, this study integrated proteomic expression data from a currently circulating epidemic strain into a metabolic model of *B. pertussis* to identify major changes that occur in metabolism between the biofilm and planktonic states. To our knowledge this is the first extensive study into specific metabolic pathways within biofilms of *B. pertussis*. Although utilised metabolic pathways differ from species to species, many of the changes in this study have been identified and confirmed experimentally in other species reinforcing the strength of the metabolic models. The major metabolic differences identified in this study relate to the TCA cycle, amino acid metabolism and virulence.

The TCA cycle is the major central metabolic pathway for many aerobic organisms, therefore it was surprising to find that there was altered flux for the TCA cycle. There was a major shift in planktonic cells to utilise the glyoxylate shunt instead of completing the full TCA cycle. The TCA reactions with increased flux in biofilms leads to the production of NADH, NADPH and ATP. Variation in the glyoxylate shunt has been observed in the biofilms of other species. When the glyoxylate shunt was disabled in *Pseudomonas aeruginosa*, there was an increase in biofilm formation (59), which is reflected in this study. It was suggested that the increased extracellular polymeric substances (EPS) produced when the glyoxylate shunt was disabled could lead to higher survival in the microaerobic conditions of the cystic fibrosis lung environment (59). Additionally, there was an increased level of glyoxylate activity in *Candida albicans* cells dispersed from biofilm (60). It is hypothesised that as the glyoxylate shunt is activated to increase nutrient versatility, it may be an anticipatory reaction for low nutrient levels while searching for a new colonisation location (60). The dispersed cells may reflect a planktonic lifestyle while established mature biofilm cells may utilise the network of cells to share nutrients and hence have a decreased requirement of the glyoxylate shunt. Targeting the TCA cycle has been suggested as a potential therapeutic strategy against biofilms (17, 61). Additional models that were created using *B. pertussis* biofilm and planktonic proteomic expression data from de Gouw *et al*. (14) had similar differences in the TCA cycle to our results (Supplementary Figure S2).

Increased polyhydroxybutyrate (PHB) and superoxide dismutase activity may explain the increased survivability of *B. pertussis* biofilm cells. Reactions regarding PHB synthesis were identified as unique in the biofilm model and superoxide dismutase activity was increased in the biofilm model. While most reactions surrounding acetyl-CoA were downregulated in the biofilm model or unique in the planktonic model, a set of reactions that involve the conversion of acetyl-CoA to PHB were unique in the biofilm models. The PHB reactions lead to an increase in cytoplasmic PHB demand. It has been previously reported that cytoplasmic PHB inclusions exist in *B. pertussis* cells (41). These inclusions are generated when there are high levels of carbon in the environment or when *B. pertussis* cells are undergoing iron starvation (41, 62). It has been hypothesised that the cells generate PHB inclusions as an energy reserve for when cells are in harsh environments (41). As the PHB demand reactions are increased in the biofilm cells, this may lead to increased cell survivability that has been observed in *B. pertussis* biofilms (15). Furthermore, an increase in the superoxide dismutase activity is protective as it leads to a decrease in superoxide which, as a free radical, may lead to cellular damage (63, 64). Put together, these two factors may be a reason for increased cell survivability of *B. pertussis* biofilm cells and may lead to persistent infections.

Many pathways for glycerophospholipid metabolism were seen to be active in both metabolic models. Lipid remodelling can occur when cells are exposed to harsh environments or antimicrobials (65). The changes in the lipids have been linked to increased survivability and persistence in pathogenic bacterial biofilms (66). The models in this study suggest that there would be differing levels of certain glycerophospholipids in the periplasm between planktonic and biofilm cells. Of particular interest is cardiolipin, identified as a unique lipid synthesis pathway in the planktonic model. Previous studies have shown that cardiolipin depletion led to decreased biofilm in *E. coli* (67). Cardiolipins have been identified in *B. pertussis* previously however, studies have not remarked on biofilm cells (68). It is important to note that the proteomic data captures enzymes that are active at a particular time point rather than a measurement of lipids, therefore, the changes identified in these reactions merely provide a snapshot of the time from which the proteins were harvested. The planktonic cells may continuously synthesise cardiolipin while mature biofilm cells either synthesise cardiolipin at an earlier stage of biofilm or have a lower requirement for cardiolipin. To further investigate the changes in lipid composition, future lipidomic analysis of biofilm cells would be necessary. Nevertheless, the evidence from the metabolic models in this study suggest differing lipid composition in biofilm and planktonic *B. pertussis* cell membranes. Another interesting change that was linked to the cardiolipin pathway was the exportation of glycerol out of the cell. While the planktonic cells exported glycerol, the biofilm cells exported glycerol 3-phosphate. The reason for this is unexplored and it would be interesting to identify whether varying concentrations of glycerol 3-phosphate or glycerol would have a regulatory role in *B. pertussis* (69).

Many regulatory processes are involved in *B. pertussis* biofilm formation (70). This study identified multiple amino acid processing pathways that varied between biofilm and planktonic cells. It has been reported previously that amino acids and their intermediates can act as regulatory molecules (71–73). The planktonic cells showed increased arginine biosynthesis and aspartate transaminase activity. It has been shown in other species that arginine may increase biofilm formation although higher concentrations have an inhibitory effect (72) and increase antibiotic killing (74). In addition, arginine may disrupt bacterial coaggregation to prevent biofilm formation (75). Finally, the arginine biosynthesis process leads to urea by-products which have been previously shown to disrupt the EPS matrix of biofilms (76). The evidence in these studies combined with results from this present study points to a tightly regulated arginine synthesis pathway which is partially fed by the aspartate transaminase reaction. While the planktonic model pushed aspartate to the arginine biosynthesis pathway, the biofilm cells pushed aspartate to the threonine degradation pathway and towards the synthesis of N-carbamoyl-L-aspartate (through to orotate) leading to an overall reduction of available arginine. The orotate was uniquely exported out of the biofilm cells rather than processed for pyrimidine metabolism. Orotate therefore has the potential to act as a signalling molecule for *B. pertussis* biofilm cells. The pathways regarding carbamoyl phosphate synthesis which leads to orotate synthesis has been shown to be altered in biofilm formation in other species (77, 78). This orotate production pathway was specific to both the biofilm model in this study and the biofilm model created from the data of de Gouw *et al*. (14), reinforcing the possible importance of orotate for *B. pertussis* biofilms.

Within the process to synthesise orotate is the incorporation of aspartate to carbamoyl phosphate (Figure 7A). Aspartate conversion from oxaloacetate and GLU (aspartate transaminase reaction) was seen to be a downregulated pathway in both biofilm metabolic models compared to their planktonic counterparts. Aspartate has been seen to inhibit biofilm formation in multiple *Staphylococcus* species (79). Furthermore, the same specific aspartate transaminase reaction was downregulated in *Streptococcus pneumoniae* biofilms (80). Thus, it is likely that aspartate may also have an inhibitory effect on *B. pertussis* biofilms.

Finally, there was a change in the processing of alanine and valine between the biofilm and planktonic models. The models suggest that β-alanine is important for planktonic cells while L-alanine is more vital for biofilm cells. The biofilm model had unique reactions transporting L-alanine into the periplasm. A recent study showed that alanine metabolism activity displays a unique pattern based on spatial distribution of cells within a heterogenous biofilm (81). The alanine metabolism changes identified in the present study may relate to a particular layer of the biofilm. It would be interesting to see the specific active reactions between the layers of biofilm. Proteomic analysis on layers of biofilm have previously been performed using laser ablation sample transfer (LAST) and label free quantification mass spectrometry (82). A combination of LAST and the methods used in this study may be able to confirm the changes in alanine metabolism. Furthermore, valine was uniquely exported out of the biofilm model. This confirms what has been observed in other Gram-negative species that had increased valine secretion in biofilm conditions (83). It was suggested that within the low oxygen and reduced growth rate conditions of biofilms, there would be an excess of pyruvate leading to an increase in valine and alanine biosynthesis (83). The discussed changes in amino acid metabolism pathways are biofilm specific and therefore it is possible that the secreted molecules could work as signalling molecules potentially triggering biofilm formation.

Major expression differences were also identified for virulence factors. Prn was found to be downregulated in biofilm cells. This is contrasted by previous results that have found an increase in Prn in *B. pertussis* biofilm cells (14, 16–18). However, it should be noted that de Gouw *et al*. had slightly decreased levels of Prn in biofilm cells compared to mid-log planktonic cells while there was a small upregulation compared to stationary phase cells (14). As this current study compared with log phase cells, the difference in Prn may be related to variations in planktonic phases of growth. Nevertheless, to our knowledge, this is the first report of a major downregulation of Prn in *B. pertussis* biofilm cells. As one of the three ACV components, the change identified supports the hypothesis that the planktonic based ACV may not protect as efficiently against cells in the biofilm condition. The utility of biofilm related proteins as novel vaccine antigens has been demonstrated previously (14, 16, 84). This study provides additional targets that may be explored further in addition to identifying key metabolic pathways that may be crucial to disrupting the biofilm lifestyle.

The virulence factor, adenylate cyclase toxin, CyaA, was upregulated in the biofilm cells. CyaA has been shown to interact with FHA and decrease biofilm formation in a concentration dependent manner (43). Furthermore, it was found that the addition of exogenous CyaA can lead to the diffusion of preformed *B. pertussis* biofilms (43). It has been shown that beyond 96 h of incubation the rate of *B. pertussis* biofilm formation can plateau (18). It is possible that *B. pertussis* utilises the activity of CyaA to diffuse cells from the biofilm to control growth in addition to allowing the spread of biofilm cells.

The Bvg system has been shown to regulate most of the virulence factors in *B. pertussis* (85). This study identified increased expression of the BvgA regulator and BvgS in biofilms while the expression of BvgR was similar between the two conditions (Figure 2B and Table 1). Upregulation of BipA underscores the importance of the Bvg^i^ phase in the biofilm existence (14, 86). However, it should be noted that there were also changes in both Bvg^+^ and Bvg^-^ protein regulation. Unexpectedly, there was a wide variability in protein expression within Bvg regulated proteins. For example, in addition to varied levels between CyaA and Prn mentioned above, Dnt was upregulated while many proteins of the type III secretion system (T3SS) were downregulated. Within both the upregulated and downregulated proteins, 13-15% of the proteins were Bvg^+^. Furthermore, there was a significantly higher proportion (14%) of proteins upregulated in biofilms that were Bvg^-^ compared to the proportion (4%) of proteins downregulated that were Bvg^-^ (Fishers exact test, *p* = 0.0002). These findings suggest that there may be a heterogeneous mixture of Bvg mode cells distributed throughout the biofilm. As there are variations in the nutrients diffused throughout a biofilm (87, 88), it is rational to expect that phenotypic modulation would occur based on the nutrients available within the microenvironment for each given cell.

It has been shown that there are varying levels of diffusion of CO_2_ and O_2_ within a biofilm (88–90). *B. pertussis* encodes regulatory systems (PlrSR and *Bpe*GReg) that respond to fluctuations in concentrations of CO_2_ and O_2_ (91, 92). While *Bpe*GReg was not identified in this study, the PlrSR system was found to be upregulated in the biofilm cells and may be indicative of varying levels of CO_2_ within the biofilm. Furthermore, the PlrSR system may lead to the change in the expression of Bvg proteins measured in this study (91). There was a range of additional response and transcriptional regulators with differential expression in biofilm cells (Supplementary Table S2). The findings in this study point to other major regulatory systems that affect the regulation of virulence proteins in biofilm conditions.

Traditional analysis of proteomic data involves grouping proteins into broad functional categories to identify major changes, but it lacks the precision to examine specific changes in metabolism. The incorporation of the proteomic data in the GSMM in this study provided new insights into the metabolic pathways that were altered between biofilm and planktonic cells. The use of GSMMs offers a different method to analyse proteomic data by viewing the connected network in a holistic way. The metabolic modelling method is designed to utilise the expression data solely as cues for particular pathways and therefore inaccuracies in measurements may have been reduced by mapping the data to a metabolic network and linking the proteins between reactions. Further studies confirming the changes in metabolism would increase confidence in the results identified. Nevertheless, an initial glimpse into the active metabolic pathways within *B. pertussis* biofilm cells was established. Parallels with other organisms that have similar fluctuations in biofilm pathways reinforce value of models and help validate the results identified by the models.

It is important to note that the biofilm cells were grown in an artificial media (THIJS media) designed to optimise *B. pertussis* growth (41). Thus, the metabolic models may be reflective of growth specifically in this medium. It would be interesting to see how growth in different media such as media more representative of the respiratory environment or co-culture with epithelial cells would affect the metabolic models (93). Furthermore, in addition to spatial analysis, it would be worthwhile to asses temporal metabolic models to identify the changes that occur throughout biofilm development (94). Finally, planktonic cells in the log phase were used for the comparison, there may be more similarities between the two conditions when planktonic cells are grown for a longer period in stationary phase.

In conclusion, this study compared the proteomic expression of biofilm and planktonic *B. pertussis* cells and identified key changes between the conditions including an upregulation of toxins (adenylate cyclase toxin and dermonecrotic toxin) and downregulation of pertactin and type III secretion system proteins in biofilm cells. Incorporation of proteomic data into a genome scale metabolic model demonstrated major metabolic changes that occur during biofilm conditions in *B. pertussis*. Notably, it was found that biofilm cells utilised the full TCA cycle while planktonic cells pushed flux through the glyoxylate shunt. There was an increase in PHB accumulation and superoxide dismutase activity which may lead to increased persistence of biofilm cells. Valine was uniquely exported of out of biofilm cells, which may have a role in cell-to-cell communication. Our study highlights the utility of integrated expression data into metabolic modelling. Overall, the changes identified in this study helps lay the groundwork for further studies into *B. pertussis* biofilms and its role in pathogenesis.

## Authors’ contributions

H.S., L.D.W.L., and R.L. designed the study. L.D.W.L., L.Z., M.J.R. and R.L. provided critical analysis and discussion, H.S. wrote the first draft and all authors contributed to the final manuscript.

## References

1. Galanis, E., A. S. King, P. Varughese, and S. A. Halperin. 2006. Changing epidemiology and emerging risk groups for pertussis. Can. Med. Assoc. J. 174: 451–452.

2. de Melker, H. E., M. A. Conyn-van Spaendonck, H. C. Rümke, J. K. van Wijngaarden, F. R. Mooi, and J. F. Schellekens. 1997. Pertussis in The Netherlands: an outbreak despite high levels of immunization with whole-cell vaccine. Emerg. Infect. Dis. 3: 175–178.

3. Güriş, D., P. M. Strebel, B. Bardenheier, M. Brennan, R. Tachdjian, E. Finch, M. Wharton, and J. R. Livengood. 1999. Changing epidemiology of pertussis in the United States: increasing reported incidence among adolescents and adults, 1990-1996. Clin. Infect. Dis. 28: 1230–1237.

4. Campbell, P. T., J. M. McCaw, P. McIntyre, and J. McVernon. 2015. Defining long-term drivers of pertussis resurgence, and optimal vaccine control strategies. Vaccine 33: 5794–5800.

5. Octavia, S., R. P. Maharjan, V. Sintchenko, G. Stevenson, P. R. Reeves, G. L. Gilbert, and R. Lan. 2011. Insight into Evolution of *Bordetella pertussis* from Comparative Genomic Analysis: Evidence of Vaccine-Driven Selection. Mol. Biol. Evol. 28: 707–715.

6. Mooi, F. R., N. A. Van Der Maas, and H. E. De Melker. 2014. Pertussis resurgence: waning immunity and pathogen adaptation -two sides of the same coin. Epidemiol. Infect. 142: 685–694.

7. Conover, M. S., G. P. Sloan, C. F. Love, N. Sukumar, and R. Deora. 2010. The Bps polysaccharide of *Bordetella pertussis* promotes colonization and biofilm formation in the nose by functioning as an adhesin. Mol. Microbiol. 77: 1439–1455.

8. Soane, M. C., A. Jackson, D. Maskell, A. Allen, P. Keig, A. Dewar, G. Dougan, and R. Wilson. 2000. Interaction of *Bordetella pertussis* with human respiratory mucosa in vitro. Respir. Med. 94: 791–799.

9. Serra, D. O., M. S. Conover, L. Arnal, G. P. Sloan, M. E. Rodriguez, O. M. Yantorno, and R. Deora. 2011. FHA-mediated cell-substrate and cell-cell adhesions are critical for *Bordetella pertussis* biofilm formation on abiotic surfaces and in the mouse nose and the trachea. PLoS One 6: e28811.

10. Paddock, C. D., G. N. Sanden, J. D. Cherry, A. A. Gal, C. Langston, K. M. Tatti, K.-H. Wu, C. S. Goldsmith, P. W. Greer, J. L. Montague, M. T. Eliason, R. C. Holman, J. Guarner, W.-J. Shieh, and S. R. Zaki. 2008. Pathology and Pathogenesis of Fatal *Bordetella pertussis* Infection in Infants. Clin. Infect. Dis. 47: 328–338.

11. Cattelan, N., J. Jennings-Gee, P. Dubey, O. M. Yantorno, and R. Deora. 2017. Hyperbiofilm Formation by *Bordetella pertussis* Strains Correlates with Enhanced Virulence Traits. Infect. Immun. 85: e00373–00317.

12. Conover, M. S., M. Mishra, and R. Deora. 2011. Extracellular DNA is essential for maintaining *Bordetella* biofilm integrity on abiotic surfaces and in the upper respiratory tract of mice. PLoS One 6: e16861–e16861.

13. Mishra, M., G. Parise, K. D. Jackson, D. J. Wozniak, and R. Deora. 2005. The BvgAS Signal Transduction System Regulates Biofilm Development in Bordetella. J. Bacteriol. 187: 1474.

14. de Gouw, D., D. O. Serra, M. I. de Jonge, P. W. Hermans, H. J. Wessels, A. Zomer, O. M. Yantorno, D. A. Diavatopoulos, and F. R. Mooi. 2014. The vaccine potential of *Bordetella pertussis* biofilm-derived membrane proteins. Emerg. Microbes Infect. 3: e58.

15. Dorji, D., R. M. Graham, P. Richmond, A. Keil, and T. K. Mukkur. 2016. Biofilm forming potential and antimicrobial susceptibility of newly emerged Western Australian *Bordetella pertussis* clinical isolates. Biofouling 32: 1141–1152.

16. Carriquiriborde, F., P. Martin Aispuro, N. Ambrosis, E. Zurita, D. Bottero, M. E. Gaillard, C. Castuma, E. Rudi, A. Lodeiro, and D. F. Hozbor. 2021. Pertussis Vaccine Candidate Based on Outer Membrane Vesicles Derived From Biofilm Culture. Front. Immunol. 12: 730434.

17. Arnal, L., T. Grunert, N. Cattelan, D. de Gouw, M. I. Villalba, D. O. Serra, F. R. Mooi, M. Ehling-Schulz, and O. M. Yantorno. 2015. *Bordetella pertussis* Isolates from Argentinean Whooping Cough Patients Display Enhanced Biofilm Formation Capacity Compared to Tohama I Reference Strain. Front. Microbiol. 6: 1352.

18. Serra, D. O., G. Lucking, F. Weiland, S. Schulz, A. Gorg, O. M. Yantorno, and M. Ehling-Schulz. 2008. Proteome approaches combined with Fourier transform infrared spectroscopy revealed a distinctive biofilm physiology in Bordetella pertussis. Proteomics 8: 4995–5010.

19. Moon, K., R. P. Bonocora, D. D. Kim, Q. Chen, J. T. Wade, S. Stibitz, and D. M. Hinton. 2017. The BvgAS Regulon of Bordetella pertussis. mBio 8: e01526–01517.

20. Belcher, T., I. MacArthur, J. D. King, G. C. Langridge, M. Mayho, J. Parkhill, and A. Preston. 2020. Fundamental differences in physiology of *Bordetella pertussis* dependent on the two-component system Bvg revealed by gene essentiality studies. Microb. Genom. 6: mgen000496.

21. Nicholson, T. L., M. S. Conover, and R. Deora. 2012. Transcriptome Profiling Reveals Stage-Specific Production and Requirement of Flagella during Biofilm Development in *Bordetella bronchiseptica*. PLoS One 7: e49166.

22. Irie, Y., S. Mattoo, and M. H. Yuk. 2004. The Bvg Virulence Control System Regulates Biofilm Formation in *Bordetella bronchiseptica*. J. Bacteriol. 186: 5692.

23. Sisti, F., D.-G. Ha apos, G. A. Toole, D. Hozbor, and J. Fernández. 2013. Cyclic-di-GMP signalling regulates motility and biofilm formation in *Bordetella bronchiseptica*. Microbiology 159: 869–879.

24. Orth, J. D., I. Thiele, and B. Ø. Palsson. 2010. What is flux balance analysis? Nat. Biotechnol. 28: 245–248.

25. Fong, N. L., J. A. Lerman, I. Lam, B. O. Palsson, and P. Charusanti. 2013. Reconciling a *Salmonella enterica* metabolic model with experimental data confirms that overexpression of the glyoxylate shunt can rescue a lethal ppc deletion mutant. FEMS Microbiol. Lett. 342: 62–69.

26. Metz, Z. P., T. Ding, and D. J. Baumler. 2018. Using genome-scale metabolic models to compare serovars of the foodborne pathogen Listeria monocytogenes. PLoS One 13:e0198584.

27. Lobel, L., N. Sigal, I. Borovok, E. Ruppin, and A. A. Herskovits. 2012. Integrative Genomic Analysis Identifies Isoleucine and CodY as Regulators of *Listeria monocytogenes* Virulence. PLoS Genet. 8: e1002887.

28. Lee, D.-S., H. Burd, J. Liu, E. Almaas, O. Wiest, A.-L. Barabási, Z. N. Oltvai, and V. Kapatral. 2009. Comparative Genome-Scale Metabolic Reconstruction and Flux Balance Analysis of Multiple *Staphylococcus aureus* Genomes Identify Novel Antimicrobial Drug Targets. J. Bacteriol. 191: 4015–4024.

29. Rienksma, R. A., P. J. Schaap, V. A. P. Martins dos Santos, and M. Suarez-Diez. 2018. Modeling the Metabolic State of *Mycobacterium tuberculosis* Upon Infection. Front. Cell. Infect. Microbiol. 8: 264.

30. Gu, C., G. B. Kim, W. J. Kim, H. U. Kim, and S. Y. Lee. 2019. Current status and applications of genome-scale metabolic models. Genome Biol. 20: 121.

31. Branco Dos Santos, F., B. G. Olivier, J. Boele, V. Smessaert, P. De Rop, P. Krumpochova, G. W. Klau, M. Giera, P. Dehottay, B. Teusink, and P. Goffin. 2017. Probing the Genome-Scale Metabolic Landscape of *Bordetella pertussis*, the Causative Agent of Whooping Cough. Appl. Environ. Microbiol. 83: e01528–01517.

32. Fyson, N., J. King, T. Belcher, A. Preston, and C. Colijn. 2017. A curated genome-scale metabolic model of *Bordetella pertussis* metabolism. PLoS Comput. Biol. 13: e1005639.

33. Gonyar, L. A., P. E. Gelbach, D. G. McDuffie, A. F. Koeppel, Q. Chen, G. Lee, L. M. Temple, S. Stibitz, E. L. Hewlett, J. A. Papin, F. H. Damron, and J. C. Eby. 2019. *In Vivo* Gene Essentiality and Metabolism in *Bordetella pertussis*. mSphere 4: e00694–00618.

34. Åkesson, M., J. Förster, and J. Nielsen. 2004. Integration of gene expression data into genome-scale metabolic models. Metab. Eng. 6: 285–293.

35. Zur, H., E. Ruppin, and T. Shlomi. 2010. iMAT: an integrative metabolic analysis tool. Bioinformatics 26: 3140–3142.

36. Stempler, S., K. Yizhak, and E. Ruppin. 2014. Integrating Transcriptomics with Metabolic Modeling Predicts Biomarkers and Drug Targets for Alzheimer’s Disease. PLoS One 9: e105383.

37. Safarchi, A., S. Octavia, S. Z. Wu, S. Kaur, V. Sintchenko, G. L. Gilbert, N. Wood, P. McIntyre, H. Marshall, A. D. Keil, and R. Lan. 2016. Genomic dissection of Australian *Bordetella pertussis* isolates from the 2008-2012 epidemic. J. Infect. 72: 468–477.

38. Safarchi, A., S. Octavia, L. D. W. Luu, C. Y. Tay, V. Sintchenko, N. Wood, H. Marshall, P. McIntyre, and R. Lan. 2016. Better colonisation of newly emerged *Bordetella pertussis* in the co-infection mouse model study. Vaccine 34: 3967–3971.

39. Safarchi, A., S. Octavia, L. D. W. Luu, C. Y. Tay, V. Sintchenko, N. Wood, H. Marshall, P. McIntyre, and R. Lan. 2015. Pertactin negative *Bordetella pertussis* demonstrates higher fitness under vaccine selection pressure in a mixed infection model. Vaccine 33: 6277–6281.

40. Luu, L. D. W., S. Octavia, L. Zhong, M. J. Raftery, V. Sintchenko, and R. Lan. 2018. Proteomic Adaptation of Australian Epidemic Bordetella pertussis. Proteomics 18: 1700237.

41. Thalen, M., I. J. van den, W. Jiskoot, B. Zomer, P. Roholl, C. de Gooijer, C. Beuvery, and J. Tramper. 1999. Rational medium design for *Bordetella pertussis*: basic metabolism. J. Biotechnol. 75: 147–159.

42. Luu, L. D. W., S. Octavia, L. Zhong, M. Raftery, V. Sintchenko, and R. Lan. 2017. Characterisation of the *Bordetella pertussis* secretome under different media. J. Proteomics 158: 43–51.

43. Hoffman, C., J. Eby, M. Gray, F. Heath Damron, J. Melvin, P. Cotter, and E. Hewlett. 2017. *Bordetella* adenylate cyclase toxin interacts with filamentous haemagglutinin to inhibit biofilm formation in vitro. Mol. Microbiol. 103: 214–228.

44. Bjerkan, G., E. Witsø, and K. Bergh. 2009. Sonication is superior to scraping for retrieval of bacteria in biofilm on titanium and steel surfaces in vitro. Acta Orthop. 80: 245–250.

45. Noorian, P., J. Hu, Z. Chen, S. Kjelleberg, M. R. Wilkins, S. Sun, and D. McDougald. 2017. Pyomelanin produced by *Vibrio cholerae* confers resistance to predation by Acanthamoeba castellanii. FEMS Microbiol. Ecol. 93.

46. Cattelan, N., O. M. Yantorno, and R. Deora. 2018. Structural Analysis of *Bordetella pertussis* Biofilms by Confocal Laser Scanning Microscopy. Bio Protocol 8: e2953.

47. Heydorn, A., A. T. Nielsen, M. Hentzer, C. Sternberg, M. Givskov, B. K. Ersboll, and S. Molin. 2000. Quantification of biofilm structures by the novel computer program COMSTAT. Microbiology 146 (Pt 10): 2395–2407.

48. Storey, J. D., and R. Tibshirani. 2003. Statistical significance for genomewide studies. Proc. Natl. Acad. Sci. U.S.A. 100: 9440–9445.

49. Bart, M. J., S. R. Harris, A. Advani, Y. Arakawa, D. Bottero, V. Bouchez, P. K. Cassiday, C.-S. Chiang, T. Dalby, N. K. Fry, M. E. Gaillard, M. van Gent, N. Guiso, H. O. Hallander, E. T. Harvill, Q. He, H. G. J. van der Heide, K. Heuvelman, D. F. Hozbor, K. Kamachi, G. I. Karataev, R. Lan, A. Lutyńska, R. P. Maharjan, J. Mertsola, T. Miyamura, S. Octavia, A. Preston, M. A. Quail, V. Sintchenko, P. Stefanelli, M. L. Tondella, R. S. W. Tsang, Y. Xu, S.-M. Yao, S. Zhang, J. Parkhill, and F. R. Mooi. 2014. Global Population Structure and Evolution of *Bordetella pertussis* and Their Relationship with Vaccination. mBio 5: e01074–01014.

50. Perez-Riverol, Y., J. Bai, C. Bandla, D. García-Seisdedos, S. Hewapathirana, S. Kamatchinathan, D. J. Kundu, A. Prakash, A. Frericks-Zipper, M. Eisenacher, M. Walzer, S. Wang, A. Brazma, and J. A. Vizcaíno. 2022. The PRIDE database resources in 2022: a hub for mass spectrometry-based proteomics evidences. Nucleic Acids Res. 50: D543–d552.

51. Heirendt, L., S. Arreckx, T. Pfau, S. N. Mendoza, A. Richelle, A. Heinken, H. S. Haraldsdóttir, J. Wachowiak, S. M. Keating, V. Vlasov, S. Magnusdóttir, C. Y. Ng, G. Preciat, A. Žagare, S. H. J. Chan, M. K. Aurich, C. M. Clancy, J. Modamio, J. T. Sauls, A. Noronha, A. Bordbar, B. Cousins, D. C. El Assal, L. V. Valcarcel, I. Apaolaza, S. Ghaderi, M. Ahookhosh, M. Ben Guebila, A. Kostromins, N. Sompairac, H. M. Le, D. Ma, Y. Sun, L. Wang, J. T. Yurkovich, M. A. P. Oliveira, P. T. Vuong, L. P. El Assal, I. Kuperstein, A. Zinovyev, H. S. Hinton, W. A. Bryant, F. J. Aragón Artacho, F. J. Planes, E. Stalidzans, A. Maass, S. Vempala, M. Hucka, M. A. Saunders, C. D. Maranas, N. E. Lewis, T. Sauter, B. Ø. Palsson, I. Thiele, and R. M. T. Fleming. 2019. Creation and analysis of biochemical constraint-based models using the COBRA Toolbox v.3.0. Nat. Protoc. 14: 639–702.

52. Feist, A. M., C. S. Henry, J. L. Reed, M. Krummenacker, A. R. Joyce, P. D. Karp, L. J. Broadbelt, V. Hatzimanikatis, and B. Ø. Palsson. 2007. A genome-scale metabolic reconstruction for *Escherichia coli* K-12 MG1655 that accounts for 1260 ORFs and thermodynamic information. Mol. Syst. Biol. 3: 121–121.

53. Izac, M., D. Garnier, D. Speck, and N. D. Lindley. 2015. A Functional Tricarboxylic Acid Cycle Operates during Growth of *Bordetella pertussis* on Amino Acid Mixtures as Sole Carbon Substrates. PLoS One 10: e0145251.

54. Malik-Sheriff, R. S., M. Glont, T. V. N. Nguyen, K. Tiwari, M. G. Roberts, A. Xavier, M. T. Vu, J. Men, M. Maire, S. Kananathan, E. L. Fairbanks, J. P. Meyer, C. Arankalle, T. M. Varusai, V. Knight-Schrijver, L. Li, C. Dueñas-Roca, G. Dass, S. M. Keating, Y. M. Park, N. Buso, N. Rodriguez, M. Hucka, and H. Hermjakob. 2019. BioModels—15 years of sharing computational models in life science. Nucleic Acids Res. 48: D407–D415.

55. Mahadevan, R., and C. H. Schilling. 2003. The effects of alternate optimal solutions in constraint-based genome-scale metabolic models. Metab. Eng. 5: 264–276.

56. Montezano, D., L. Meek, R. Gupta, L. E. Bermudez, and J. C. M. Bermudez. 2015. Flux Balance Analysis with Objective Function Defined by Proteomics Data— Metabolism of *Mycobacterium tuberculosis* Exposed to Mefloquine. PLoS One 10: e0134014.

57. Goto, M., A. Abe, T. Hanawa, and A. Kuwae. 2021. Bcr4 is a Chaperone for the Inner Rod Protein in the *Bordetella* Type III Secretion System. bioRxiv: 2021.2009.2028.462275.

58. Bosch, A., D. Serra, C. Prieto, J. Schmitt, D. Naumann, and O. Yantorno. 2005. Characterization of *Bordetella pertussis* growing as biofilm by chemical analysis and FT-IR spectroscopy. Applied Microbiology and Biotechnology 71: 736.

59. Ahn, S., J. Jung, I.-A. Jang, E. L. Madsen, and W. Park. 2016. Role of Glyoxylate Shunt in Oxidative Stress Response. J. Biol. Chem. 291: 11928–11938.

60. Uppuluri, P., M. Acosta Zaldívar, M. Z. Anderson, M. J. Dunn, J. Berman, J. L. Lopez Ribot, and J. R. Köhler. 2018. *Candida albicans* Dispersed Cells Are Developmentally Distinct from Biofilm and Planktonic Cells. mBio 9: e01338–01318.

61. Yahya, M., U. Hamid, M. Norfatimah, and R. Kambol. 2014. *In silico* analysis of essential tricarboxylic acid cycle enzymes from biofilm-forming bacteria. Trends in Bioinformatics 7: 19–26.

62. Alvarez Hayes, J., Y. Lamberti, K. Surmann, F. Schmidt, U. Völker, and M. E. Rodriguez. 2015. Shotgun proteome analysis of *Bordetella pertussis* reveals a distinct influence of iron availability on the bacterial metabolism, virulence, and defense response. Proteomics 15: 2258–2266.

63. Suo, Y., Y. Huang, Y. Liu, C. Shi, and X. Shi. 2012. The expression of superoxide dismutase (SOD) and a putative ABC transporter permease is inversely correlated during biofilm formation in *Listeria monocytogenes* 4b G. PLoS One 7: e48467–e48467.

64. De Groote, M. A., U. A. Ochsner, M. U. Shiloh, C. Nathan, J. M. McCord, M. C. Dinauer, S. J. Libby, A. Vazquez-Torres, Y. Xu, and F. C. Fang. 1997. Periplasmic superoxide dismutase protects *Salmonella* from products of phagocyte NADPH-oxidase and nitric oxide□synthase. Proc. Natl. Acad. Sci. U.S.A. 94: 13997–14001.

65. Seydlova, G., J. Beranova, I. Bibova, A. Dienstbier, J. Drzmisek, J. Masin, R. Fiser, I. Konopasek, and B. Vecerek. 2017. The extent of the temperature-induced membrane remodeling in two closely related *Bordetella* species reflects their adaptation to diverse environmental niches. J. Biol. Chem. 292: 8048–8058.

66. Dubois-Brissonnet, F., E. Trotier, and R. Briandet. 2016. The Biofilm Lifestyle Involves an Increase in Bacterial Membrane Saturated Fatty Acids. Front. Microbiol. 7: 1673.

67. Nepper, J. F., Y. C. Lin, and D. B. Weibel. 2019. Rcs Phosphorelay Activation in Cardiolipin-Deficient *Escherichia coli* Reduces Biofilm Formation. J. Bacteriol. 201: e00804–00818.

68. Kawai, Y., and A. Moribayashi. 1982. Characteristic lipids of *Bordetella pertussis:* simple fatty acid composition, hydroxy fatty acids, and an ornithine-containing lipid. J. Bacteriol. 151: 996–1005.

69. Wapnir, R. A., and L. Stiel. 1985. Regulation of gluconeogenesis by glycerol and its phosphorylated derivatives. Biochem. Med. 33: 141–148.

70. Cattelan, N., P. Dubey, L. Arnal, O. M. Yantorno, and R. Deora. 2016. *Bordetella* biofilms: a lifestyle leading to persistent infections. Pathog. Dis. 74: ftv108.

71. Cava, F., H. Lam, M. A. de Pedro, and M. K. Waldor. 2011. Emerging knowledge of regulatory roles of D-amino acids in bacteria. Cell Mol. Life Sci. 68: 817–831.

72. Goh, S.-N., A. Fernandez, S.-Z. Ang, W.-Y. Lau, D.-L. Ng, and E. S. G. Cheah. 2013. Effects of Different Amino Acids on Biofilm Growth, Swimming Motility and Twitching Motility in *Escherichia coli* BL21. Journal of Biology and Life Science 4: 13.

73. Aliashkevich, A., L. Alvarez, and F. Cava. 2018. New Insights Into the Mechanisms and Biological Roles of D-Amino Acids in Complex Eco-Systems. Front. Microbiol. 9: 683.

74. Borriello, G., L. Richards, G. D. Ehrlich, and P. S. Stewart. 2006. Arginine or Nitrate Enhances Antibiotic Susceptibility of *Pseudomonas aeruginosa* in Biofilms. Antimicrob. Agents Chemother. 50: 382–384.

75. Kolderman, E., D. Bettampadi, D. Samarian, S. E. Dowd, B. Foxman, N. S. Jakubovics, and A. H. Rickard. 2015. L-Arginine Destabilizes Oral Multi-Species Biofilm Communities Developed in Human Saliva. PLoS One 10: e0121835.

76. Brindle, E. R., D. A. Miller, and P. S. Stewart. 2011. Hydrodynamic deformation and removal of *Staphylococcus epidermidis* biofilms treated with urea, chlorhexidine, iron chloride, or DispersinB. Biotechnol. Bioeng. 108: 2968–2977.

77. Zhuo, T., W. Rou, X. Song, J. Guo, X. Fan, G. G. Kamau, and H. Zou. 2015. Molecular study on the *carAB* operon reveals that *carB* gene is required for swimming and biofilm formation in *Xanthomonas citri* subsp. citri. BMC Microbiol. 15: 225–225.

78. Pisithkul, T., J. W. Schroeder, E. A. Trujillo, P. Yeesin, D. M. Stevenson, T. Chaiamarit, J. J. Coon, J. D. Wang, D. Amador-Noguez, J.-M. Ghigo, and E. P. Greenberg. 2019. Metabolic Remodeling during Biofilm Development of Bacillus subtilis. mBio 10: e00623–00619.

79. Yang, H., M. Wang, J. Yu, and H. Wei. 2015. Aspartate inhibits *Staphylococcus aureus* biofilm formation. FEMS Microbiol. Lett. 362: fnv025.

80. Allan, R. N., P. Skipp, J. Jefferies, S. C. Clarke, S. N. Faust, L. Hall-Stoodley, and J. Webb. 2014. Pronounced metabolic changes in adaptation to biofilm growth by *Streptococcus pneumoniae*. PLoS One 9: e107015.

81. Díaz-Pascual, F., M. Lempp, K. Nosho, H. Jeckel, J. K. Jo, K. Neuhaus, R. Hartmann, E. Jelli, M. F. Hansen, A. Price-Whelan, L. E. P. Dietrich, H. Link, and K. Drescher. 2021. Spatial alanine metabolism determines local growth dynamics of *Escherichia coli* colonies. eLife 10: e70794.

82. Pulukkody, A. C., Y. P. Yung, F. Donnarumma, K. K. Murray, R. P. Carlson, and L. Hanley. 2021. Spatially resolved analysis of *Pseudomonas aeruginosa* biofilm proteomes measured by laser ablation sample transfer. PLoS One 16: e0250911.

83. Valle, J., S. Da Re, S. Schmid, D. Skurnik, R. D’Ari, and J. M. Ghigo. 2008. The amino acid valine is secreted in continuous-flow bacterial biofilms. J. Bacteriol. 190: 264–274.

84. Dorji, D., R. M. Graham, A. K. Singh, J. P. Ramsay, P. Price, and S. Lee. 2019. Immunogenicity and protective potential of *Bordetella pertussis* biofilm and its associated antigens in a murine model. Cell. Immunol. 337: 42–47.

85. Cotter, P. A., and A. M. Jones. 2003. Phosphorelay control of virulence gene expression in *Bordetella*. Trends Microbiol. 11: 367–373.

86. Deora, R. 2004. Multiple mechanisms of *bipA* gene regulation by the *Bordetella* BvgAS phosphorelay system. Trends Microbiol. 12: 63–65.

87. Salgar-Chaparro, S. J., K. Lepkova, T. Pojtanabuntoeng, A. Darwin, L. L. Machuca, and A. J. M. Stams. 2020. Nutrient Level Determines Biofilm Characteristics and Subsequent Impact on Microbial Corrosion and Biocide Effectiveness. Appl. Environ. Microbiol. 86: e02885–02819.

88. Stewart, P. S. 2003. Diffusion in biofilms. J. Bacteriol. 185: 1485–1491.

89. Stewart, P. S., T. Zhang, R. Xu, B. Pitts, M. C. Walters, F. Roe, J. Kikhney, and A. Moter. 2016. Reaction–diffusion theory explains hypoxia and heterogeneous growth within microbial biofilms associated with chronic infections. npj Biofilms Microbomes 2: 16012.

90. Patel, T. D., and T. R. Bott. 1991. Oxygen diffusion through a developing biofilm of Pseudomonas fluorescens. J. Chem. Technol. Biotechnol. 52: 187–199.

91. Bone, M. A., A. J. Wilk, A. I. Perault, S. A. Marlatt, E. V. Scheller, R. Anthouard, Q. Chen, S. Stibitz, P. A. Cotter, and S. M. Julio. 2017. *Bordetella* PlrSR regulatory system controls BvgAS activity and virulence in the lower respiratory tract. Proc. Natl. Acad. Sci. U.S.A. 114: E1519–E1527.

92. Wan, X., J. R. Tuckerman, J. A. Saito, T. A. K. Freitas, J. S. Newhouse, J. R. Denery, M. Y. Galperin, G. Gonzalez, M.-A. Gilles-Gonzalez, and M. Alam. 2009. Globins Synthesize the Second Messenger Bis-(3’–5’)-Cyclic Diguanosine Monophosphate in Bacteria. J. Mol. Biol. 388: 262–270.

93. Palmer, K. L., L. M. Aye, and M. Whiteley. 2007. Nutritional cues control *Pseudomonas aeruginosa* multicellular behavior in cystic fibrosis sputum. J. Bacteriol. 189: 8079–8087.

94. Siriwach, R., F. Matsuda, K. Yano, and M. Y. Hirai. 2020. Drought Stress Responses in Context-Specific Genome-Scale Metabolic Models of Arabidopsis thaliana. Metabolites 10: 159.

