## Supplementary Figure S1, Supplementary Figure S2, Supplementary Table S1 for "Integrating proteomic data with metabolic modelling provides insight into key pathways of *Bordetella pertussis* biofilms"

Supplementary figures and tables

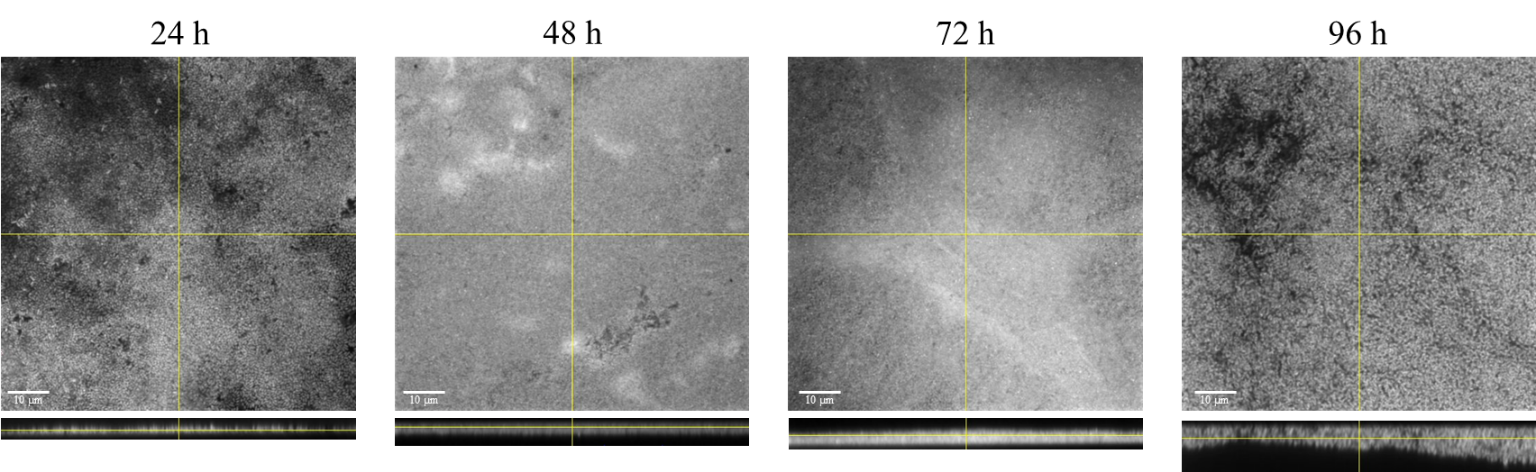

**B**

**A**

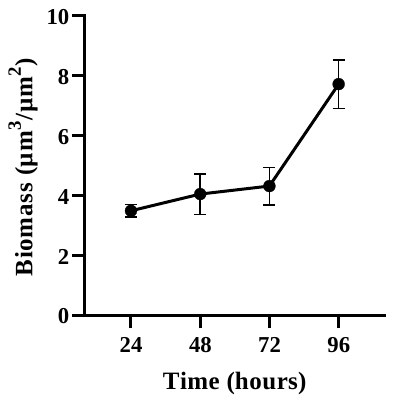

**Supplementary Figure S1** **A** Confocal laser scanning microscopy micrographs of currently circulating strain, L1423 *B. pertussis* biofilm formation from 24 h to 96 h. Biofilms were grown on glass cover slides imaged in a z-stack setting. Images are presented in xy and xz planes. Images are represented in colourblind-safe grayscale (1) **B** Biomass calculated at each time point using the COMSTAT2 ImageJ plugin. Three biological replicates per time point were performed with 3 randomly selected field of views for extracted values.

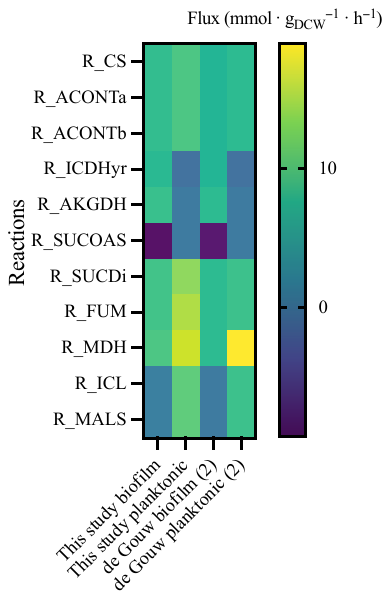

**Supplementary Figure S2 Flux values for reactions in the tricarboxylic acid cycle between additional models.** Reactions extracted from FBA results for the models in this study (biofilm and planktonic) and de Gouw *et al.* (biofilm and planktonic) (2). R_CS, type II citrate synthase; R_ACONTa, citrate hydrolase; R_ACONTb, aconitate hydratase; R_ICDHyr, isocitrate dehydrogenase; R_AKGDH, α-ketoglutarate dehydrogenase; R_SUCOAS, succinyl-CoA synthetase; R_SUCDi, succinate dehydrogenase; R_FUM, fumarate hydratase; R_MDH, malate dehydrogenase; R_ICL, isocitrate lyase; R_MALS, malate synthase.

**Strain differences between de Gouw *et al.* (2) study and this study**

The strains used by de Gouw *et al*. (2) and this study are both of the same genotype *ptxP3-ptxA1-prn2*. The *ptxP3* strains have overtaken the previously dominant *ptxP1* strains and have been linked to the resurgence in pertussis cases (3). However, there are some differences between the strains. B1917 was isolated from a Dutch patient in 2000 while L1423 was isolated in Australia during the 2008 – 2012 epidemic (4). The genomes of the strains have been previously compared and found to have 11 non-synonymous SNPs and 6 indels between them (5). These variations may account for the differences identified between the two studies. Furthermore, there are differences in the growth conditions that are described in **Supplementary Table S1**.

**Supplementary Table S1** Comparison of conditions used in the other studies from which additional iMAT models were extracted. Included are the number of reactions, metabolites and genes in each metabolic model.

|  | This study biofilm | This study planktonic | de Gouw biofilm (2) | de Gouw planktonic (2) |
| --- | --- | --- | --- | --- |
| Media | THIJS | THIJS | THIJS | THIJS |
| Incubation time | 96 h | 12 h | 72 h | 17 h/40 h |
| Strain | L1423 | L1423 | B1917 | B1917 |
| Vessel | 24 well plate | 20 mL Bioreactor tube | Polypropylene beads | 500 mL Erlenmeyer flasks |
| Media refreshment | 5 hr | NA | 5 hr then every 24 hr | NA |
| **iMAT models** |  |  |  |  |
| Reactions | 206 | 219 | 194 | 134 |
| Metabolites | 198 | 213 | 191 | 127 |
| Genes | 188 | 195 | 195 | 150 |

**Supplementary references**

1.	Johnson, J. 2012. Not seeing is not believing: improving the visibility of your fluorescence images. *Mol. Biol. Cell* 23: 754-757.

2.	de Gouw, D., D. O. Serra, M. I. de Jonge, P. W. Hermans, H. J. Wessels, A. Zomer, O. M. Yantorno, D. A. Diavatopoulos, and F. R. Mooi. 2014. The vaccine potential of *Bordetella pertussis* biofilm-derived membrane proteins. *Emerg Microbes Infect* 3: e58.

3.	Mooi, F. R., I. H. van Loo, M. van Gent, Q. He, M. J. Bart, K. J. Heuvelman, S. C. de Greeff, D. Diavatopoulos, P. Teunis, N. Nagelkerke, and J. Mertsola. 2009. *Bordetella pertussis* strains with increased toxin production associated with pertussis resurgence. *Emerg. Infect. Dis.* 15: 1206-1213.

4.	Safarchi, A., S. Octavia, S. Z. Wu, S. Kaur, V. Sintchenko, G. L. Gilbert, N. Wood, P. McIntyre, H. Marshall, A. D. Keil, and R. Lan. 2016. Genomic dissection of Australian *Bordetella pertussis* isolates from the 2008–2012 epidemic. *J. Infect.* 72: 468-477.

5.	Luu, L. D. W. 2018. Comparative proteomic analysis of Australian epidemic *Bordetella pertussis*. In *School of Biotechnology and Biomolecular Sciences*. University of New South Wales. PhD thesis
